# The GTPase activating protein Gyp6 binds Retromer and inactivates Rab7/Ypt7 to coordinate the formation of endosomal carriers

**DOI:** 10.64898/2026.01.23.701266

**Authors:** Ana Catarina Alves, Kai-En Chen, Lydie Michaillat Mayer, Qian Guo, Ella J. Stephens, Meihan Liu, Michael D. Healy, Brett M. Collins, Andreas Mayer

**Author notes:** These authors contributed equally (Ana Catarina Alves, Kai-En Chen, Brett M. Collins, Andreas Mayer).

## Abstract

The Retromer coat is conserved in all eukaryotes and is crucial for the correct intracellular sorting of many transmembrane receptors and lysosomal hydrolases. Retromer is an effector of the late endosomal small GTPase RAB7 and is also implicated in its inactivation required for proper endosomal maturation. Here, we explore the role of controlled GTP hydrolysis by the RAB7 ortholog Ypt7 in the formation of Retromer-coated membrane carriers in yeast. Proximity labelling and genetic ablation identify the GTPase Activating Protein (GAP) Gyp6 as a critical regulator of Ypt7 activity in the context or Retromer. Structural studies show that Retromer recruits Gyp6 through its Vps29 subunit, which recognises a specific PL motif and a secondary binding site in the C-terminal domain of Gyp6. This interaction does not occur with other yeast GAPs. Ablation of the Gyp6-Retromer interface or the catalytic activity of Gyp6 leads to the accumulation of tubular structures on endo-lysosomal compartments and to increased association of Ypt7 with Retromer and its cargo Vps10. These results support a model in which Gyp6 controls the switch from Ypt7-dependent Retromer coat assembly and cargo collection to the departure of the carrier through membrane fission and uncoating.

## Introduction

The delivery of transmembrane protein and lipid cargos to specific cellular compartments is an essential process in all eukaryotes and is regulated by a complex network of trafficking pathways. Collectively, these pathways guarantee the distribution of such cargoes to a great variety of specific destinations, which is vital for cellular homeostasis. Disruption or dysregulation of the cargo-trafficking system in humans is associated with various diseases, including neurodegenerative disorders such as Parkinson’s and Alzheimer’s disease (Panicker *et al*, 2021; Qureshi *et al*, 2020), and infection by bacterial and viral pathogens (Yong *et al*, 2021).

Among the key players on endo-lysosomal compartments is Retromer, a conserved complex of the proteins Vps35, Vps26 and Vps29. Retromer orchestrates the recruitment of cargoes into tubulovesicular carriers, ensuring their escape from the lysosomal degradation system and, thus, their reuse (Seaman, 2012; Gopaldass *et al*, 2024; Yong *et al*, 2022; Chen *et al*, 2019). The molecular stages governing carrier formation involve cargo recognition, Retromer oligomerization, membrane tubulation, membrane fission, and, eventually, uncoating. Recruitment of Retromer to the endosomal membrane precedes or occurs concomitantly with the selection of cargo proteins, requiring adaptors from the sorting nexin (SNX) protein family for cargo recruitment. In yeast, Retromer can form a pentameric assembly with a dimer of Vps5–Vps17, two SNX proteins with bin/amphiphysin/rvs (BAR) domains (Kovtun *et al*, 2018; Chen *et al*, 2025; Gopaldass *et al*, 2023; Seaman *et al*, 1998), or a tetramer with the Snx3 adaptor (Strochlic *et al*, 2007; Purushothaman & Ungermann, 2018; Leneva *et al*, 2021). In each case, Retromer cooperates with the SNX adaptor to bind cargo sequences and promote formation of endosomal membrane tubules into which the cargo proteins are sorted. Following tubule detachment, Retromer should dissociate from the membrane so that further rounds of cargo sorting can occur.

Retromer activity is closely linked to its interaction with small GTPases, particularly RAB7, a member of the Rab family that controls the maturation and fusion of the endosomal sorting compartment with late endosomes/lysosomes (or the vacuole in yeast (Feng *et al*, 1995; Wichmann *et al*, 1992). Rab GTPases function as molecular switches that alternate between an active GTP-bound state and an inactive GDP-bound state, and regulation of their activity is crucial for maintaining the homeostasis of the late endocytic pathway (Homma *et al*, 2021; Pylypenko *et al*, 2018; Stenmark, 2009; Kummel *et al*, 2023). This Rab switching is controlled by guanine nucleotide exchange factors (GEFs) that swap GDP for GTP to activate the Rab, and by GTPase-activating proteins (GAPs) that enhance GTP hydrolysis and Rab inactivation. The mammalian Retromer complex depends on Rab GTPases for efficient membrane recruitment (Rojas *et al*, 2008; Seaman *et al*, 2009; Modica *et al*, 2025; Priya *et al*, 2015; Harrison *et al*, 2014; Liu *et al*, 2012) and in turn impacts Rab conversion, the replacement of RAB5 by RAB7, at the endosomal compartment (Xie *et al*, 2020; Distefano *et al*, 2018; Jimenez-Orgaz *et al*, 2018; Jia *et al*, 2016; Antón-Plágaro *et al*, 2025). Retromer associates with endosomes partly through its interaction with GTP-bound RAB7. Recruitment of the GAP TBC1D5 promotes RAB7 GTP hydrolysis and inactivation. Blocking this process entails accumulation of active RAB7 and defects in endosomal maturation, which led to the suggestion that GTP-hydrolysis by RAB7 might trigger coat disassembly (Jia *et al*, 2016). However, attempts to manipulate the GAP-stimulated GTP hydrolysis by RAB7 produced inconsistent effects on localisation of endosomal cargoes such as CI-MPR, GLUT1, integrin-α5 and the HPV protein L2 (Seaman *et al*, 2009; Jimenez-Orgaz *et al*, 2018; Xie *et al*, 2020), challenging this idea. In yeast, Retromer associates with the RAB7 orthologue Ypt7 (Balderhaar *et al*, 2010; Liu *et al*, 2012; Purushothaman *et al*, 2017; Wu *et al*, 2021), but no Retromer-associated GAPs have yet been reported. Ypt7 is activated by the Mon1-Ccz1 GEF complex (Nordmann *et al*, 2010), and the yeast GAP Gyp7 regulates Ypt7 activity (Füllbrunn *et al*, 2024; Brett *et al*, 2008; Vollmer *et al*, 1999), although its specificity for RAB substrates is relatively broad *in vitro* (Vollmer *et al*, 1999). Yeast has eleven RABs and eight GAPs, the latter being characterised by a Tre/Bub2/Cdc16 (TBC) domain with a catalytic arginine-glutamine finger. Whether GAPs other than Gyp7 can control Ypt7 activity, and whether any of these might cooperate with Retromer in an analogous way to mammalian TBC1D5, is unknown.

In this study, we used a TurboID-based proteomics approach to identify yeast proteins associated with Ypt7 in its native environment. Combining the results with *in silico* modelling, structural and biochemical studies we show that Gyp6 is the only yeast GAP protein capable of associating with both Retromer and Ypt7 and elucidate its dual mechanism of binding to Retromer. We analysed its impact on Vps10 cargo recycling *in vivo*, on the formation of endosomal carriers and on Retromer coat complex formation, leading us to a model in which Gyp6 exerts spatiotemporal control over the transition from cargo collection to carrier departure.

## Results

RAB7 and its yeast homolog Ypt7 are important for the stable membrane recruitment of Retromer as well as proper maturation of the early endosome into degradative vacuolar and lysosomal compartments (Rojas *et al*, 2008; Seaman *et al*, 2009; Balderhaar *et al*, 2010; Liu *et al*, 2012; Priya *et al*, 2015; Harrison *et al*, 2014; Modica *et al*, 2025; Wu *et al*, 2021). We aimed to examine the role of GTP hydrolysis by Ypt7 in the formation of endosomal carriers in yeast to explore whether Ypt7 might constitute a GTP-triggered switch between different stages of endosomal carrier formation.

### Proximity labelling identifies Gyp3 and Gyp6 as potential GAP(s) of Ypt7

In its multiple endo-lysosomal functions, Ypt7 interacts with different partners (Numrich *et al*, 2015; Kucharczyk *et al*, 2001; Balderhaar *et al*, 2010; Cabrera *et al*, 2009). These may be regulated by the actions of different GAPs, but assigning a specific GAP to a specific Ypt7-dependent process is complicated by their potential redundancy (Fukuda, 2011). We therefore investigated the protein-protein interactions with Ypt7 by proximity labelling, which can capture transient and weak GAP-Ypt7 interactions in the native cellular environment. TurboID (a yeast optimized biotin ligase) and a V5 epitope were fused to the *Saccharomyces cerevisiae* (sc) Ypt7 N-terminus and integrated into the URA3 locus of a *ypt7*Δ mutant (**Figure 1A**) (Kim *et al*, 2014; Branon *et al*, 2018). Since expressing this fusion construct from an endogenous YPT7 promoter did not completely rescue the vacuole fragmentation phenotype of a *ypt7*Δ mutant, we expressed TurboID-V5-YPT7 (wild-type or variants) from a tetracycline-inducible promoter. This construct resulted in slightly increased Ypt7 expression levels compared to a wild-type strain (**Supplementary Fig. S1**) and completely rescued the vacuole morphology of *ypt7*Δ cells. As a control, we expressed non-fused Turbo-ID-V5 in the URA3 locus of wildtype cells.

**Figure 1:**
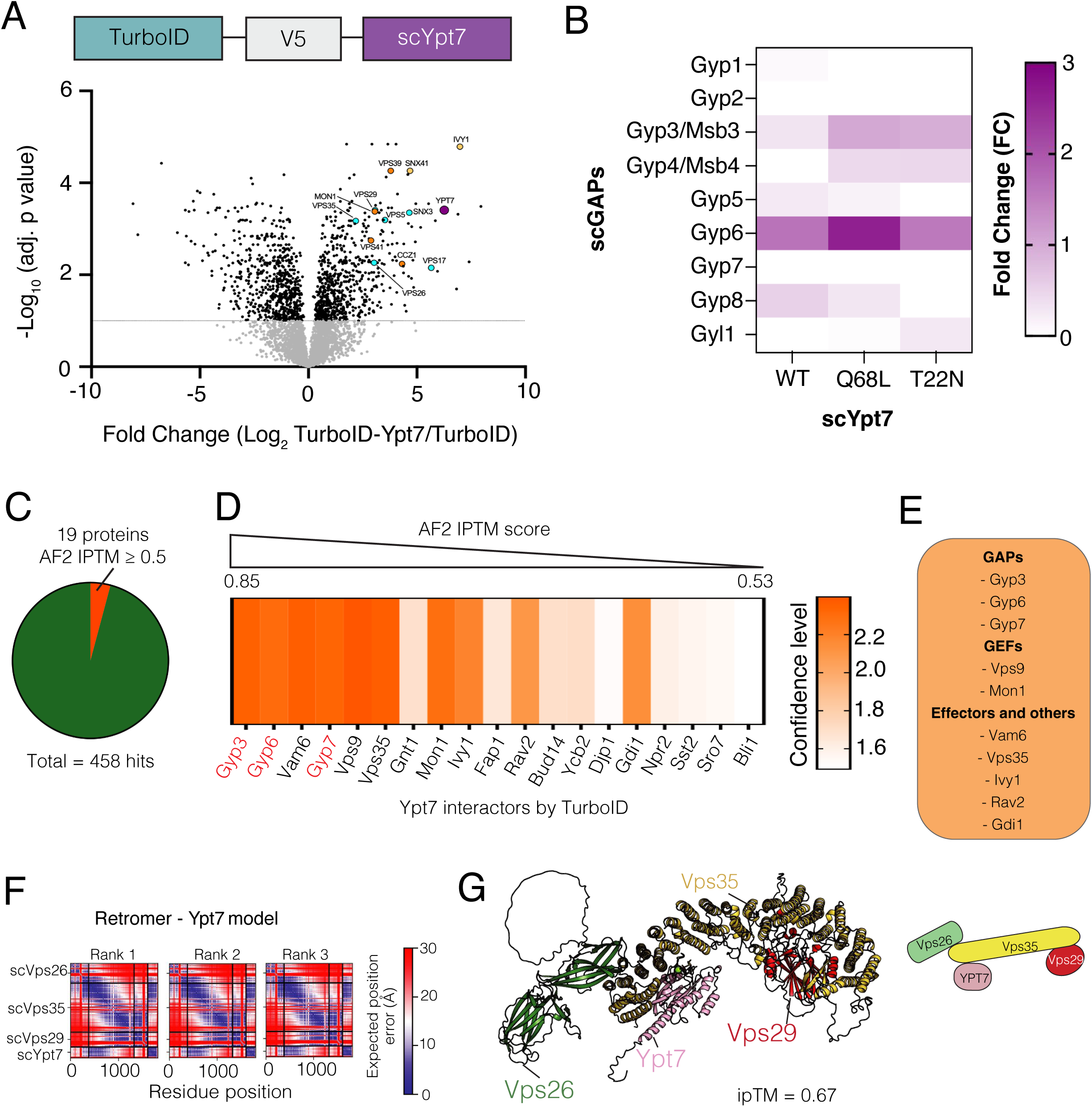
Identification of cognate GAPs of Ypt7 by proximity labelling. **(A)** Volcano plot of TurboID-Ypt7 proximity labelling versus control cells expressing only cytosolic TurboID, from a proteomics analysis of streptavidin-biotin pulldowns. Selected relevant hits are indicated and marked with color. Gray dots correspond to selected non-significant hits (significance < 0.05). All interactors detected for each Ypt7 variant are provided in the supplementary datasheet XX. **(B)** The heatmap depicts the Fold Change (FC) value of all GAP proteins for each Ypt7 form (Wild-type, Q68L, and T22N). **(C)** Schematic diagram showing the number of targets identified from the TurboID-Ypt7 with an AF2 IPTM score ≥ 0.5. **(D)** Screening of potential Ypt7 direct interactors using AF2 and AF3. The confidence of interactors was assessed based on the sum of the interfacial PTM score (iPTM), averaged across three AF2 models, AF3 models and pDOCKQ score. From left to right, the list of Ypt7 interactors is ordered from high (0.85) to low (0.53) AF2 iPTM score, respectively. **(E)** List of TurboID targets showing a high confidence score with the ability to interact directly with Ypt7. **(F)** The PAE plots for the top-three ranked models of Ypt7 – Retromer complex predicted by AlphaFold2. **(G)** Cartoon representation of the top-ranked model Ypt7 – Retromer complex. The iPTM score represents the averaged iPTM score from the three models.

We performed proximity labelling with three alleles of *YPT7*: The *YPT7* wild-type, a *ypt7^T22N^* allele coding a single amino acid substitution that interferes with GTP binding, and *ypt7^Q68L^*, which alters the switch II glutamine sidechain and reduces GTPase activity, but maintains a considerable potential for stimulating this activity by GAPs (Vollmer *et al*, 1999). *ypt7^Q68L^* was expected to stabilize Ypt7-GAP interactions while *ypt7^T22N^* is expected to impair them. Cells were labelled with biotin, lysed, and biotinylated proteins were enriched on magnetic streptavidin beads and identified by mass spectrometry. The log_2_ fold-change (FC) of peptides enriched in extracts from the wildtype TurboID-V5-YPT7 cells relative to those expressing only TurboID-V5 (control) were plotted (**Figure 1A**). Numerous known regulators and interacting partners of Ypt7 were among the best hits in TurboID-V5-Ypt7 cells, validating the approach: Mon1 and Ccz1, which form the GEF of Ypt7 (Nordmann *et al*, 2010), and Gdi1 (GDP dissociation inhibitor) (Garrett *et al*, 1994); the Retromer subunits Vps35, Vps26, and Vps29 (Seaman *et al*, 2009; Balderhaar *et al*, 2010); the HOPS subunits Vam6/Vps39 and Vps41 (acting in vacuole fusion (Seals *et al*, 2000); Ivy1 (a regulator of vacuole membrane homeostasis and nutrient sensing) (Lazar *et al*, 2002; Numrich *et al*, 2015). Along with these proteins, the Rab-GAPs Gyp6 and Gyp3/Msb3 (Albert & Gallwitz, 1999) were biotinylated by TurboID-V5-Ypt7 (**Figure 1B**). Gyp3 and Gyp6 proximity labelling increased significantly in cells carrying *ypt7^Q68L^*, probably because this variant has lower GTPase activity and hence favours interaction with GAPs. Gyp7, which has been identified as a cognate GAP of Ypt7 in previous studies (Brett *et al*, 2008; Vollmer *et al*, 1999; Stroupe, 2018), was not enriched in our Ypt7 proximity interactome.

We complemented these proteomics findings through *in silico* screening with AlphaFold2-multimer (AF2) (Mirdita *et al*, 2022). Complexes of Ypt7 with candidate proteins from our proximity labelling were predicted, and the confidence of the modelled interactions was assessed via several metrics, including the interfacial PTM (iPTM) score (**Figure 1C**), consistency of the models in AF2 and AlphaFold3 (AF3), and the pDockQ score (**Figures 1D and 1E**). With this approach, we found 19 out of 458 potential Ypt7 interactors with an iPTM score above 0.5, and 10 of these exhibited consistently high confidence scores. Most of these 19 proteins are known Ypt7 interactors (**Supplementary Fig. S2**). Among the top hits are the GAPs Gyp6, Gyp3 as well as Gyp7, which was not detected in our proximity labelling approach but was included in the computational screen due to evidence from previous studies (Brett *et al*, 2008; Vollmer *et al*, 1999; Stroupe, 2018). A high confidence interaction was also obtained for Retromer, where Ypt7 is predicted to occupy an N-terminal region on Vps35 (**Figures 1F and 1G**) (Schmid & Walter, 2025).

### Gyp6-dependent trafficking of Vps10

Since Rab-GAPs often show redundant activity, we employed two strategies to test the functional significance of Gyp3, Gyp6 and Gyp7 for Ypt7 in the context of Retromer-dependent trafficking *in vivo*. To address potential redundancy, we ablated Gyp3, Gyp6 and Gyp7 individually and in combinations in *S. cerevisiae*. Their relationship to Ypt7 was probed through the *ypt7^Q68^*^L^ allele, which has 5-fold reduced GTPase activity but can still be stimulated by GAPs (Vollmer *et al*, 1999). It can hence provide a sensitized background for probing the effect of GAPs on sorting of Retromer cargo *in vivo*. We categorized the mutants according to the fraction of cells that accumulated the Retromer cargo Vps10 fused with mNeonGreen (Vps10^mNG^) on the (FM 4-64-labeled) vacuolar membrane (**Figures 2A and 2B**). Vps10^mNG^ was detected on vacuoles in less than 2% of all YPT7 wildtype cells, and this proportion remained low in the *ypt7^Q68L^* strain (5% of all cells). Both *gyp3*Δ and *gyp6*Δ single mutants and *gyp6*Δ *gyp3*Δ double mutants were very responsive to the presence of *ypt7^Q68L^*. Introducing *ypt7^Q68L^* into *gyp6*Δ cells increased the fraction showing vacuolar Vps10^mNG^ from 10% to 35%; for *gyp6*Δ *gyp3*Δ the fraction increased from 63% to 100% (**Figures 2A and 2B**). The *gyp6*Δ *gyp3*Δ *ypt7^Q68^*^L^ cells thus phenocopied the strong sorting defect of the Retromer mutant *vps35*Δ. *gyp7*Δ cells showed no impact on Vps10^mNG^ localization, neither alone nor in all possible combinations with *gyp3*Δ*, gyp6*Δ and *ypt7*^Q68L^ mutations **(Supplementary Figures S3, S4A and S4B)**. Gyp6 also affected a phenotype that is linked to the missorting of Vps10, the secretion of the vacuolar hydrolase carboxypeptidase Y. *gyp6*Δ cells secreted significantly more carboxypeptidase Y than wildtype and *gyp3*Δ cells, and *gyp3*Δ *gyp6*Δ revealed a strong additive effect (**Figure 2C**).

**Figure 2:**
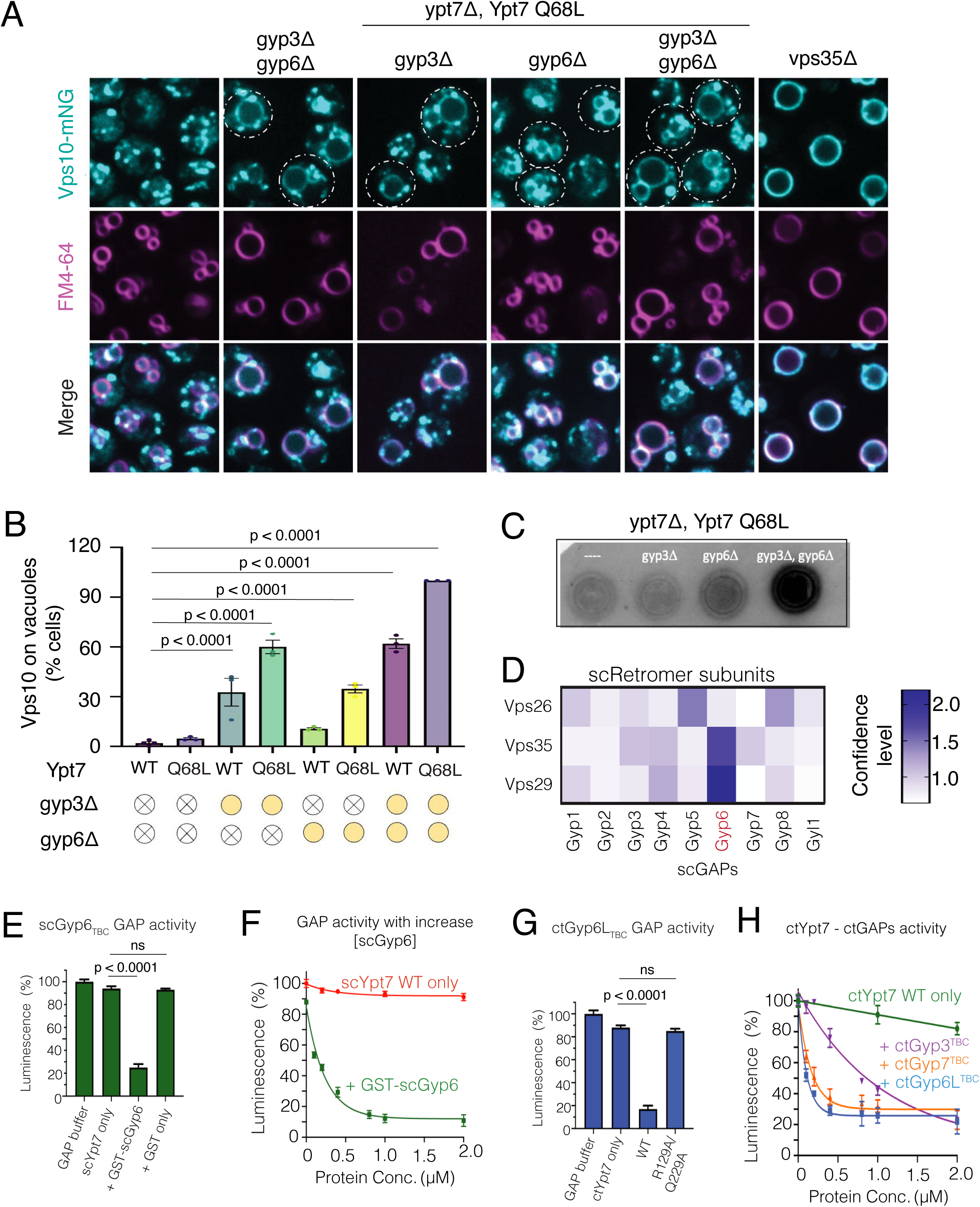
Vps10 recycling defect in GAP mutants. **(A)** Live-cell fluorescence microscopy of yeast expressing Vps10^mNeonGreen^ (cyan) following FM 4-64 pulse-chase labelling to mark the vacuolar membrane (magenta). Representative z-stack average projections from 5 single-focal planes are shown. Scale bar, 5 µm. **(B)** Quantification of Vps10 colocalization with the vacuolar membrane marker FM 4-64, obtained from three independent experiments (n=50 cells per experiment were classified). Mean ± s.e.m. are shown. *P*-values were calculated by One-way ANOVA. **(C)** Western blot analysis of carboxypeptidase Y (CPY) secretion in the indicated yeast strains. **(D)** Heat map showing the *in silico* screening of scRetromer subunits with scRab GAPs using AF2 and AF3. The confidence score of potential interactors was assessed based on the sum of the iPTM score from three AF2, AF3 models. **(E)** GAP activity of GST-tagged Gyp6 towards Ypt7. **(F)** GTP hydrolysis of Ypt7 in the presence of indicated concentration of GST-Gyp6. **(G)** Comparison of the GAP activity assay of wild-type and mutant form of ctGyp6L^TBC^ towards ctYpt7. **(H)** GTP hydrolysis of ctYpt7 in the presence of indicated concentration of GAPs including ctGyp3^TBC^ (purple line), ctGyp6L^TBC^ (blue line) or ctGyp7^TBC^ (orange line).

To further explore how these GAP family members might link to Retromer and Ypt7, we performed an *in silico* screen of all nine yeast Rab GAPs against the three individual Retromer subunits using both AF2 and AF3. The confidence of the binding models was ranked by combining iPTM scores from the AF runs with the pDockQ score as mentioned above. Among the nine *S. cerevisiae* GAPs screened, Gyp6 showed the highest predicted interaction scores, preferentially associating with Vps29 and Vps35 (**Figure 2D; Supplementary Fig. S5A-S5B**). For the interaction with Ypt7, AlphaFold models predict that Gyp6 binds to the expected surface patch of Ypt7 (**Supplementary Fig. S5C-S5D**), inserting its canonical Arginine finger (Arg155) to enable GAP activity (**Supplementary Fig. S5E**). To confirm that Gyp6 functions as a GAP for Ypt7, we assessed GTPase activity using the GTPase-Glo^TM^ assay. Here, GAP activity was tested using Gyp6 from both *S. cerevisiae* and *Chaetomium thermophilum* to assess conservation across different yeast species. Using the recombinant full-length Gyp6, we observed robust activation of Ypt7 (**Figure 2E**) with GAP activity increasing in a concentration dependent manner (**Figure 2F**). Similar *in vitro* GAP activity was also observed using the *C. thermophilum* homologues, Gyp6-like protein TBC domain (ctGyp6L^TBC^) and ctYpt7. Wild-type ctGyp6L^TBC^ strongly activated the GTPase activity of ctYpt7 (**Figure 2G**). In contrast, the ctGyp6L^TBC^ R129A/Q229A mutant, where the conserved arginine finger and the YxQ motif that are necessary for GAP activity were substituted (Pan *et al*, 2006), showed no detectable activity (**Figure 2G**). While the *C. thermophilum* GAPs ctGyp7^TBC^ and ctGyp6L^TBC^ both displayed significant GAP activity, activation by ctGyp3^TBC^ was weaker (**Figure 2H**). In combination with its strong signal in proximity labelling, its impact on Vps10 sorting, and confidently predicted interaction using AlphaFold, these observations suggest Gyp6 as the preferred GAP for Ypt7 in the context of Retromer activity. In line with this, Gyp6 also localizes to dot-like structures, where we found a Gyp6^mNG^ fusion protein to frequently colocalize with Vps35^mCherry^ in puncta (**Supplementary Fig. S6**), which are likely to represent endosomal compartments (Ali *et al*, 2004; Suda *et al*, 2013; Brunet *et al*, 2016; Thomas *et al*, 2021).

### Gyp6 interacts with Retromer via a PL-motif

Predictions of the *S. cerevisiae* Retromer–Gyp6 complex using AF3 indicate that two regions of the Retromer trimer are involved in the interaction (**Figure 3A; Supplementary Fig. S7A-S7C**). According to the models, a high confidence interaction is observed between Vps29 and a Pro-Leu (PL) motif of Gyp6, which is located within the long hinge loop (Leu48 to Leu142) next to the arginine finger of the TBC domain (**Figures 3A, 3B, 3C and 3D**). Such PL-motifs are a frequent feature of Vps29-binding proteins across evolution (Antón-Plágaro *et al*, 2025; Guo *et al*, 2024; Healy *et al*, 2023; Yao *et al*, 2018; Romano-Moreno *et al*, 2017; Barlocher *et al*, 2017; Crawley-Snowdon *et al*, 2020). This PL motif is unique to Gyp6 among yeast Rab GAPs (**Figure 3E**) but it is conserved across Gyp6 homologs in different yeast species (**Figure 3F**). Confident predictions of PL-mediated Gyp6-Retromer interactions were also obtained for the *C. thermophilum* and *S. pombe* orthologs (**Supplementary Fig. S8**). However, the PL motif in ctGyp6L and spGyp6 is located on a shorter loop between helix 4 and helix 5 of the TBC domain, in contrast to the longer loop found in *S. cerevisiae* Gyp6 (**Supplementary Fig. S8**).

**Figure 3:**
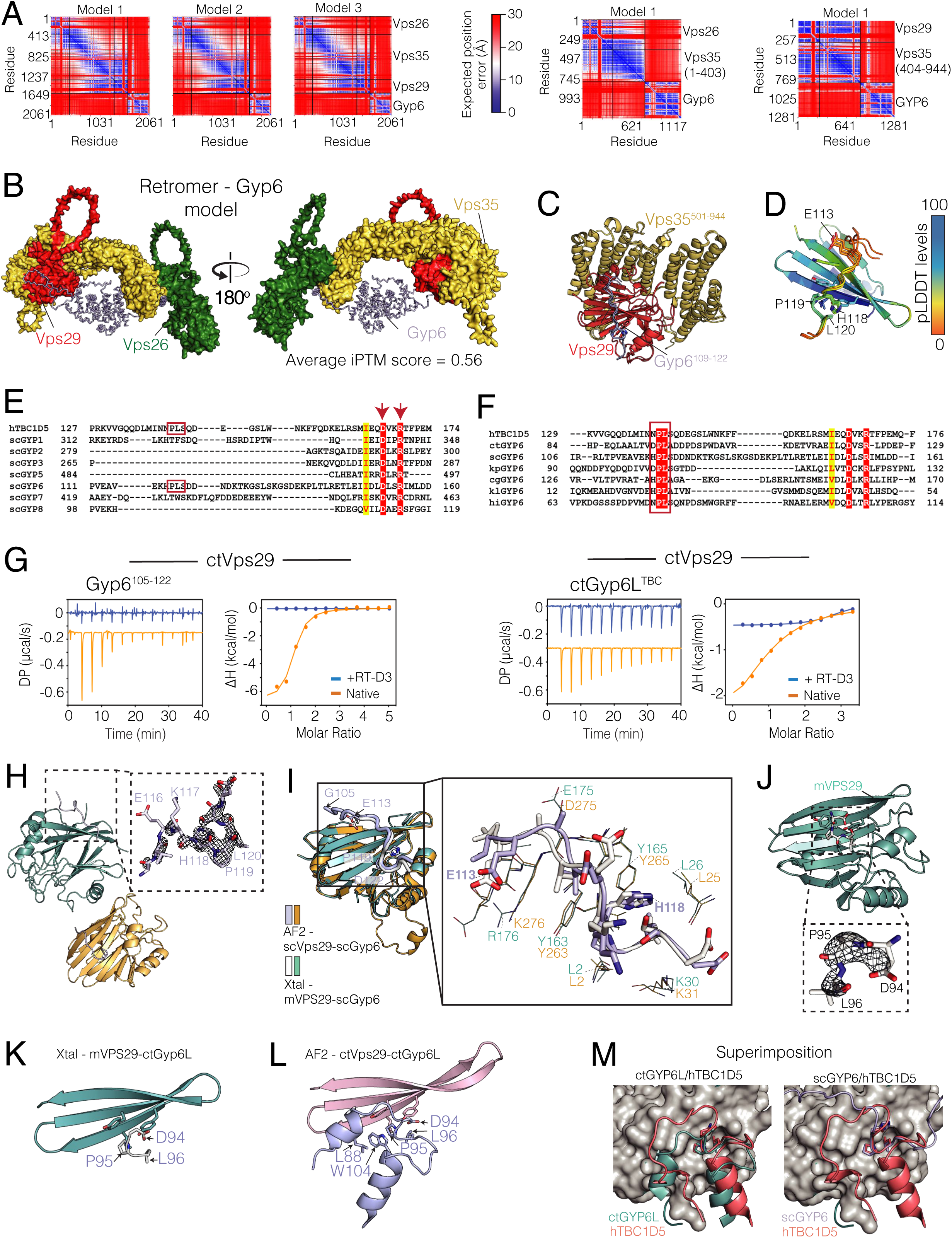
Structural basis confirming the Retromer and Gyp6 association through the PL motif. **(A)** PAE plots showing the top-ranked AF2 models of Retromer - Gyp6 complex (left), the Vps35^1-403^ – Vps26 – Gyp6 (middle) and VPS35^501-944^ – Vps29 – Gyp6 subcomplexes (right). **(B)** Structural representation of the top-ranked model of the Retromer – Gyp6 complex. For clarity, Gyp6 is shown in ribbon. **(C)** Overlay of the three predicted models of Vps35^501-944^ – Vps29 – Gyp6 complex. For clarity, only the key PL motif of Gyp6 involved in binding is shown. **(D)** Close-up view of the contact region between Vps29 and the PL motif of Gyp6. Models are coloured according to the pLDDT score. **(E, F)** Sequence alignment of yeast Gyp family proteins highlighting that the PL motif is uniquely present in Gyp6 and is conserved across Gyp6 orthologs. (**G**) ITC measurements of Gyp6^105-122^ peptide (left) and ctGyp6L^TBC^ domain (right) binding to ctVps29, in the presence or absence of Vps29-binding cyclic peptide, RT-D3. ITC graphs show the integrated and normalized data with a 1 to 1 ratio binding ratio. **(H)** Crystal structure of the mVPS29 – Gyp6^105-122^ complex, showing two molecules in the asymmetric unit. The dashed box indicates the electron density of the Gyp6^105-122^ peptide, corresponding to a stimulated-annealing OMIT Fo-Fc map contoured at 3σ. **(I)** Superimposition of the mVPS29 – Gyp6^105-122^ crystal structure (light teal) and the corresponding AF2 model (bright orange), highlighting the key binding residues (scGyp6 residues G105 to D122 shown in light purple). For clarity, the scVps29 disordered loop composed of P163 to S245 is not shown. **(J)** Crystal structure of the mVPS29 – ctGyp6L^87-107^ complex, showing a single molecule in the asymmetric unit. The dashed box highlights the electron density of the ctGyp6L^87-107^ peptide, as shown in (**H**). **(K)** Close-up view of the mVPS29 – ctGyp6L^87-107^ crystal structure and **(L)** the corresponding AF2 model as shown in Figure (**I**). **(M)** Surface representation of the ctVps29 – ctGyp6L (left) and Vps29 – Gyp6 (right) complexes, illustrating the two distinct PL motif binding modes. ctGyp6L adopts a conformation more similar to TBC1D5 (highlighted in red) than to Gyp6.

To validate the AlphaFold models, we confirmed the contribution of the PL motif to the Gyp6-Retromer interaction by isothermal calorimetry (ITC) and X-ray crystallography. Due to challenges in producing sufficient *S. cerevisiae* Retromer subunits for ITC and structural studies, we used *C. thermophilum* and human Retromer subunits for these experiments. Using ITC, we found that a Gyp6^105-122^ peptide containing the PL motif binds to ctVps29 with a *K*_d_ of 1.4 μM (**Figure 3G**; **Table 1**). Under the same condition, we found the recombinant ctGyp6L^TBC^ binds to ctVps29 with a *K*_d_ of 27 μM, but the PL motif containing peptide, ctGyp6L^87-107^ failed to interact due to the weaker affinity (**Figure 3G; Supplementary Fig. 9, Table 1**). In both cases, binding could be abolished by a known competitive Vps29 inhibitor, the cyclic peptide RT-D3 (Chen *et al*, 2021), which also contains a PL motif able to outcompete other proteins requiring the same site in Vps29 (**Figure 3G**). This confirms that the PL motif is required for the Gyp6-Vps29 interaction.

Next, we attempted to obtain the crystal structure of Vps29 in complex with Gyp6 or Gyp6-derived peptides. From multiple crystallization screens, diffraction-quality crystals were obtained using a combination of mouse VPS29 (mVPS29) in complex with *S. cerevisiae* Gyp6^105-122^ and ctGyp6L^87-107^ peptides. In the case of the Gyp6^105-122^ peptide complex, the structure was solved at 1.9 Å resolution (**Table 2**) with the peptide density from Val112 to Asp122 of Gyp6 clearly visible (**Figure 3H**). The region upstream of the PL motif (residues Val112 to His118) contributes to binding, including a charge interaction between Glu113 of Gyp6 and Lys276 of Vps29 (Arg176 in mouse VPS29) (**Figure 3I**). The AlphaFold models of the *S. cerevisiae* proteins correspond well with this crystal structure. In addition, they provide information on the N-terminal part of the Gyp6^105-122^ peptide that is not resolved in the crystal structure (**Figure 3I**). These models suggest that further residues upstream of the PL motif may be involved in binding, with Val100, Val104, Ile105, and Leu109 providing hydrophobic interactions with residues on Vps29 and in the Vps29–Vps35 interface.

The structures suggest two distinct PL motif binding modes. The ctGyp6L interaction involves a PL motif within a short loop region that binds Vps29 (**Figures 3K and 3L**). This short loop links two flanking helices that are tightly associated through hydrophobic interactions, forming a β-hairpin (**Figure 3L**). This arrangement resembles that of TBC1D5 binding to VPS29 (**Figure 3M**) (Jia *et al*, 2016). In this binding mode, the PL motif is the main region responsible for the contact and the interaction is weaker. By contrast, the interaction of *S. cerevisiae* Gyp6 with Vps29 shows a longer hinge loop in which additional residues reinforce the binding (**Figure 3M**). In the crystal structure of the isolated ctGyp6L^TBC^ domain, electron density was not observed for the loop containing the PL motif, suggesting it is flexible in the absence of Vps29 interaction (**Supplementary Fig. S10A-S10B**). Overall, our structural data show that Gyp6 from different yeast species shares a conserved PL motif to bind to Vps29, but the *S. cerevisiae* protein has additional contacts with Retromer that explain its higher binding affinity.

### The C-terminal tail of Gyp6 provides a second Retromer interaction site

Further analysis of the Gyp6 proteins and their interactions with Retromer revealed that Gyp6L from *C. thermophilum* contains a long, disordered C-terminal tail that is not present in *S. cerevisiae* Gyp6 (**Figure 4A**). In the ctRetromer–ctGyp6L complex Alphafold model, this disordered C-terminal tail (residues Gly676 to Gly773) binds to the rear side of the ctVps35 α-helical solenoid (**Figure 4B**). This interaction involves several Leu, Ile, and Phe residues within the disordered ctGyp6L tail, and is conserved among related species (**Figures 4C; Supplementary Fig. S11**). Interestingly, this resembles the interaction between human VPS35 and the LFa repeat motifs of human FAM21 (**Figure 4D**) (Guo *et al*, 2024). These LFa motifs in FAM21 have an inherently low affinity for human VPS35 (Guo *et al*, 2024), and we were similarly unable to measure a *K*d of ctRetromer binding to synthetic peptides of the ctGyp6L tail region by ITC (residues 680-698 and 760-773) (**Figure 4E**). It is possible that the multiple LFa-like motifs ctGyp6L (we refer these as ‘LFa-like’ due to the absence of clustered acidic residues) in the tail must cooperate with each other and with the canonical PL-motif interaction with Vps29 to enhance avidity for Retromer. To test this, we designed a scaffold to artificially increase the number of LFa-like sequences by fusing the C-terminal tail of ctGyp6L (residues 676 to 773, containing three LFa-like motifs) to the C-terminus of the archaeal protein Lsm alpha (LsmA). This protein assembles into highly stable homo-hexameric and heptameric ring structures (**Supplementary Fig. S12A-S12E**) (Collins *et al*, 2001). Using mass photometry, we found that the chimeric construct, LsmA-ctGyp6L^676-773^, adopts a size consistent with a hexamer (**Supplementary Fig. S12F**), indicating that the addition of ctGyp6L^676-773^ does not alter the oligomerization state of LsmA. We then performed ITC experiments of LsmA-ctGyp6L^676-773^ with ctRetromer. The chimeric protein showed enhanced binding affinity with a K_d_ of 5.8 µM (**Figure 4F**; **Table 1**). In contrast, a mutant construct with the LFa-like motifs substituted to alanine showed no detectable binding (**Figure 4F**), confirming that the C-terminal tail of ctGyp6L binds to the ctVps35 subunit of ctRetromer through its LFa-like motif.

**Figure 4:**
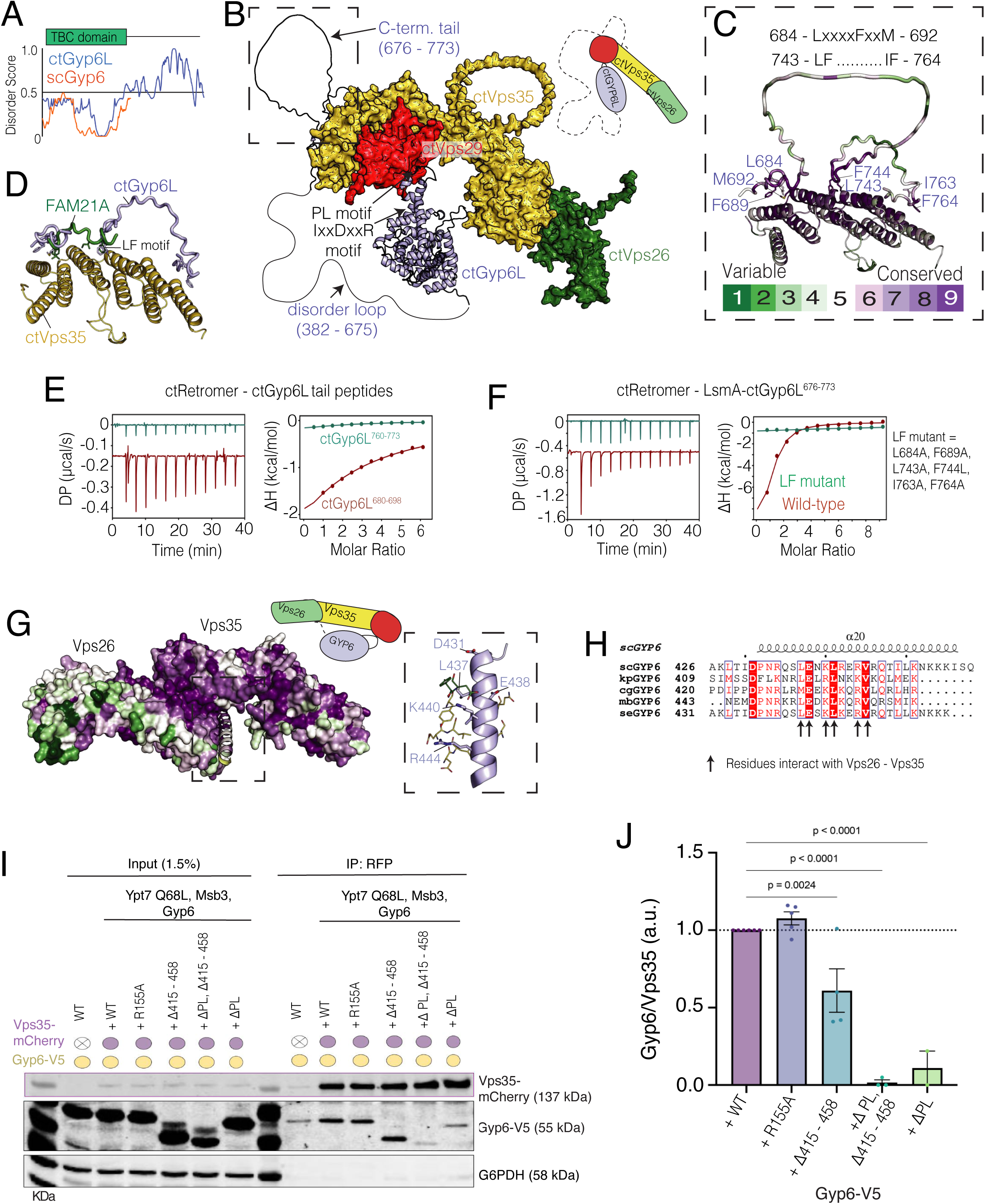
Gyp6 has a second binding site with Retromer to enhance its affinity. **(A)** Prediction of the disordered C-terminal tail of ctGyp6L and Gyp6, highlighting their differences. Disorder region predictions were mapped using the D2P2 web server. **(B)** Structural representation of the top-ranked ctRetromer – ctGyp6L model, showing the C-terminal tail of ctGyp6L binding to the rear surface of ctVps35. For clarity, ctRetromer is shown as surface and ctGyp6L as ribbon. **(C)** Sequence conservation mapped onto the ctVPS35 surface, illustrating that the LF motif of the ctGyp6L C-terminal tail (shown in ribbon) binds to ctRetromer. **(D)** Overlay of the ctGyp6L tail – ctVps35 and FAM21A LFa repeat 19 – VPS35 models, demonstrating the conservation of the LFa motifs binding model at the C-terminal end of Vps35. **(E, F)** ITC measurements of ctGyp6L^680-698^, ctGyp6L^760-733^ peptides, and LsmA-ctGyp6L^676-773^ binding to ctRetromer. All ITC graphs show the integrated and normalized data fit with a 1 to 1 ratio binding ratio. **(G)** Representation of the top-ranked AF3 model of the Retromer – Gyp6 complex, with sequence conservation mapped onto the surface. The C-terminal helix of Gyp6 binds to the Vps26 – Vps35 interface. **(H)** Sequence alignment of the Gyp6 C-terminal tail helix across the *Saccharomycetes* class. Arrows indicate conserved residues potentially involved in binding. **(I)** Co-immunoprecipitation from *ypt7^Q68L^ gyp3*Δ *gyp6*Δ *S. cerevisiae* cells. Lysates from these cells expressing the indicated variants of Gyp6^V5^ and Vps35 with or without an mCherry tag were incubated with anti-mCherry beads. The adsorbed proteins were probed by SDS-PAGE and Western blotting with the indicated antibodies. Gyp6 variants: ΔC: lacking aa 415-458; ΔPL: lacking aa 107+120. Input lanes contain 1.5% of the total lysate. **(J)** Blots from three independent experiments from (I) were quantified on a Licor Odyssey fluorescence scanner. The ratio of co-immunoprecipitated Gyp6^V5^ over Vps35^mCherry^ was normalized and set to 1 for the Gyp6 wildtype. Data represent mean ± s.e.m. from *n* = 3 independent experiments. Statistical significance was determined using one-way Anova.

In the case of *S. cerevisiae* Gyp6, the C-terminal tail is much shorter than that of *C. thermophilum* Gyp6 and is instead predicted to form an α-helix (**Figure 4A**). According to AF3 modelling, this α-helical C-terminus may bind at the Vps26–Vps35 interface through several conserved charged and hydrophobic residues (**Figures 4G-4H; Supplementary Fig. S7A**). To test this prediction, we performed co-immunoprecipitation assays from cells co-expressing Vps35^yomCherry^ and Gyp6^V5^ from their endogenous promoter. Both the wild-type and the catalytically dead variant Gyp6^R155A-V5^ bound Retromer (**Figures 4I-4J**). However, Gyp6^ΔPL-V5^ (lacking PL motif residues 105-122) and Gyp6^ΔC-V5^ (lacking the C-terminal helix residues 415-458) both showed reduced binding efficiency. Simultaneous ablation of both predicted binding sites nearly abolished Retromer binding (**Figures 4I-4J**). Overall, our data suggest that two regions of Gyp6 mediate its interaction with Retromer, one around the PL motif and the other at the C-terminal tail, where the interactions involving the C-terminal tail can vary among Gyp6 homologues.

### Gyp6 GAP activity and its binding to Retromer are required for Vps10 recycling

To examine the importance of yeast Gyp6 binding and activity for Retromer function *in vivo* we ablated or mutated the primary PL motif (Δ107-120 or R108G, L109G, T110R, V112G, P119G, L120A) and the C-terminal helix of Gyp6 (Δ415-458) and generated catalytically dead *gyp6^R155A^* (Haas *et al*, 2005). The variants were C-terminally tagged with V5 and expressed from the endogenous Gyp6 promoter in the sensitized *gyp6*Δ *gyp3*Δ *ypt7^Q68L^* background.

*gyp6*Δ *gyp3*Δ *ypt7^Q68L^* cells accumulated Vps10^mNG^ on vacuoles as previously shown (**Figure 5A**). While wild-type GYP6 partially rescued this phenotype, catalytically inactive *gyp6*^R155A^ and the variant lacking the C-terminal helix in combination with PL motif did not (**Figure 5B**). In line with this mislocalization of Vps10, which is a sorting receptor for the vacuolar hydrolase CPY (Cooper & Stevens, 1996), all three *gyp6* mutants secreted CPY into the extracellular space around yeast colonies on an agar plate, which was not the case for wild-type cells (**Figure 5C**). Thus, both Gyp6 GAP activity and its binding to Retromer are required to support Retromer function in Vps10 retrieval and vacuolar protein sorting.

**Figure 5:**
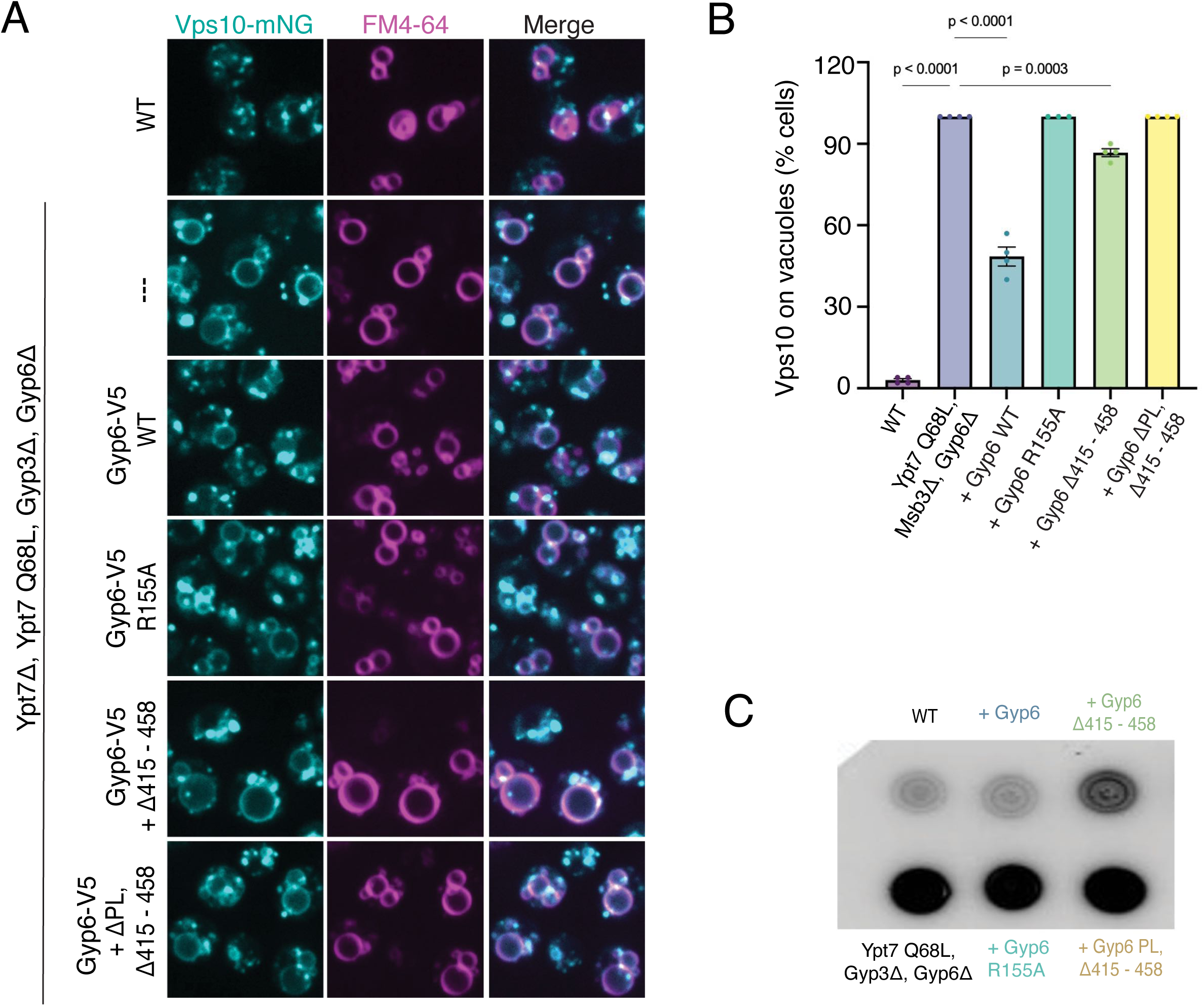
Gyp6 GAP activity and its binding to Retromer required for Vps10 recycling. **(A)** Live-cell fluorescence microscopy of yeast expressing Vps10^mNeonGreen^ (cyan) following FM 4-64 pulse-chase labelling to mark the vacuolar membrane (magenta). In Wild-type cells, Vps10 localizes to discrete puncta, largely non-overlapping with FM 4-64, consistent with its presence in late-Golgi and endosomal compartments, while it accumulates at the vacuole membrane in GAP mutants. Representative Z-stack images, from 5 single-focal planes, are shown. Scale bar, 5 μm. **(B)** Quantification of Vps10 colocalization with the vacuolar membrane marker FM 4-64, obtained from three independent experiments (n=50 cells per experiment were classified). Means and standard error of the mean are shown. *P*-values were calculated by One-way ANOVA. **(C)** Western blot analysis of carboxypeptidase Y (CPY) secretion in the indicated yeast strains. In wild-type cells, CPY is retained intracellularly, whereas secretion of CPY into the medium is observed in GAP mutants.

### Ypt7 inactivation by GAPs promotes carrier departure

Cargo retrieval requires cargo capture in sorting domains and cargo transfer into tubulovesicular structures, which then detach to form cargo carriers. Failure of Retromer-dependent transfer of Vps10 from endosomes to the Golgi leads to the accumulation of Vps10 in the vacuolar membrane (Suzuki *et al*, 2019). If this failure is generated by a complete inactivation of the sorting machinery, such as by deleting Retromer, the accumulating cargo is homogeneously distributed over the vacuolar membrane (see **Figure 2A**). The situation is different when cargo collection remains functional, but cargo exit from the compartment is limiting, e.g., due to an overload of the sorting machinery by an excess of cargo. In this case, Vps10 concentrates in small, Retromer-containing zones from which tubular structures are formed and depart (Arlt *et al*, 2015).

To differentiate which of these processes might require Gyp6 GAP activity and Ypt7 inactivation, we examined whether the GAP mutants accumulated Vps10^mNG^ cargo in dot-like structures at the vacuolar membrane (**Figure 5A**; **Figure 6A**). If these zones correspond to Retromer-dependent cargo sorting domains, Vps35^mCherry^ should co-accumulate in the same structures. In line with this, *gyp6*Δ*gyp3*Δ mutants showed larger and more intense Vps35^yomCherry^ clusters at vacuoles than wild-type cells. These Vps35 dots accumulated not only Vps10^mNG^ (**Figure 6A**) but also Ypt7 (**Figure 6B**). In *gyp6*Δ *gyp3*Δ *ypt7^Q68L^* cells, vacuolar regions concentrating Vps10, Vps35 and Ypt7 were even more prominent (**Figure 6A-6B**). Importantly, they often displayed elongated or tubular shapes that accumulate both coat components (Vps35 ^yomCherry^) and cargo (Vps10 ^mNG^). Such structures were never observed in wild-type cells. These tubular structures resemble the tubulo-vesicular vacuolar Vps10 exit zones that can be observed upon transient overexpression of Vps10, a treatment that overloads the endosomal retrieval system, generates a spill-over of Vps10 into the vacuolar membrane, and therefore triggers compensatory retrieval of the Vps10 from the vacuolar surface (Arlt *et al*, 2015). The vacuolar tubules in *gyp6*Δ *gyp3*Δ *ypt7^Q68L^* cells can hence be considered as bona fide nascent cargo carriers.

**Figure 6:**
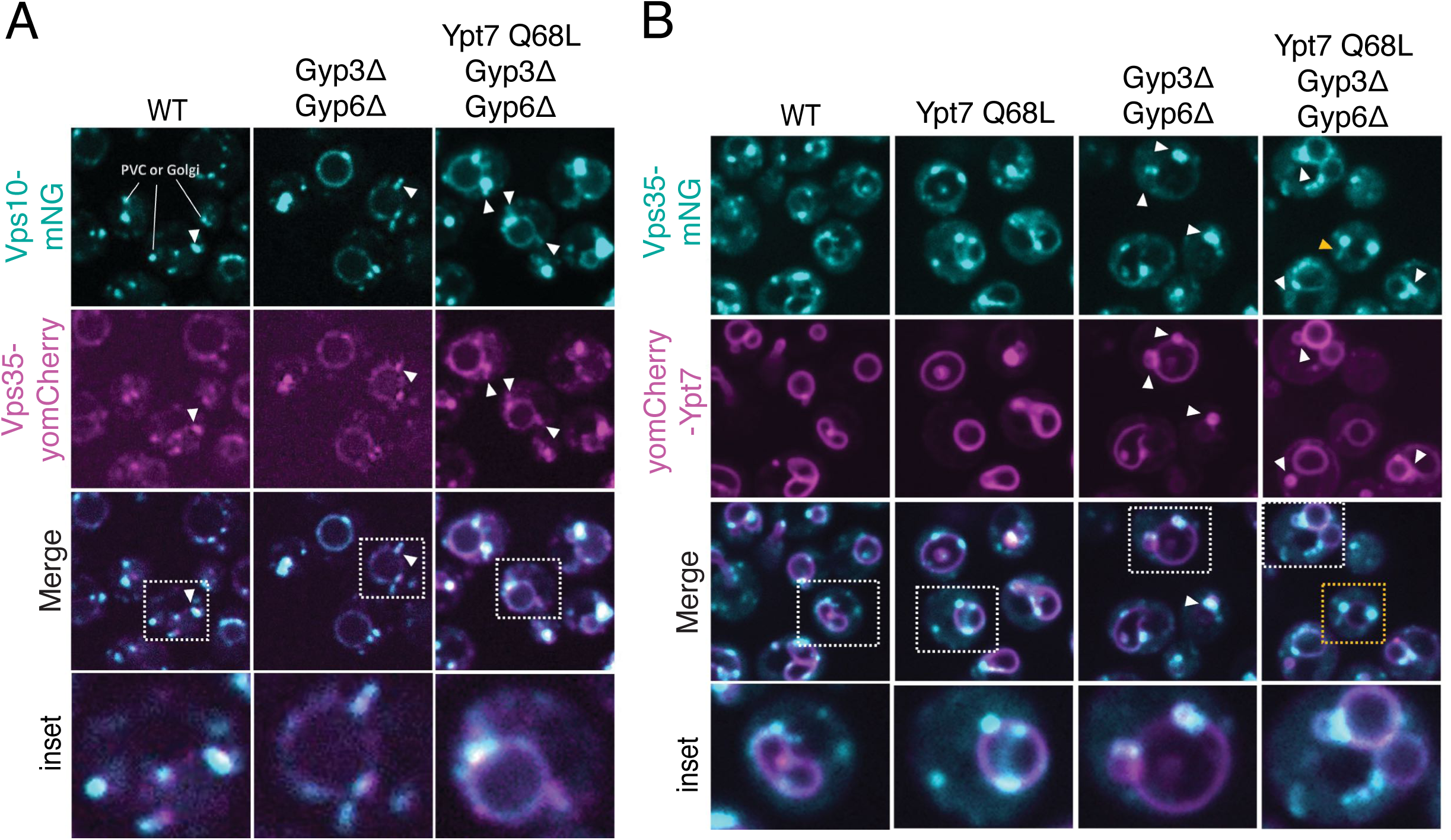
Inhibiting GAP activity leads to membrane tubule accumulation. **(A)** Live cell fluorescence microscopy of yeast cells co-expressing Vps10^mNG^ and Vps35^yomCherry^. Tubular structures that accumulate both cargo and Retromer are indicated by white arrows. Insets show the boxed areas at higher magnification. **(B)** Fluorescence microscopy of yeast cells co-expressing Vps35^mNG^ and Ypt7^yomCherry^. White arrowheads point to Vps35^mNG^ puncta co-accumulating Ypt7^yomCherry^. The yellow box and arrowhead indicate tubular structures that can be frequently observed in these mutants. Insets show the areas within the white boxes at higher magnification. Scale bars: 5 μm.

To assay whether cargo-coat complexes indeed accumulate, we performed immunoprecipitations from *gyp6*Δ*gyp3*Δ*ypt7^Q68L^* cells co-expressing Vps35^yomCherry^ and Vps10^mNG^. These cells showed two- to threefold more Vps10^mNG^ and Ypt7 associated with Vps35^yomCherry^ relative to wildtype cells (**Figure 7A**). Expression of wild-type Gyp6^V5^ in these cells attenuated this phenotype. Gyp6 variants that were catalytically dead (R155A) or lacking the PL motif and the C-terminal helix did not correct the accumulating coat-cargo-Ypt7 interactions (**Figures 7A and 7B**). Thus, Retromer-cargo complexes accumulate in the absence of Gyp6 activity or in the absence of Gyp6 recruitment to the coat.

**Figure 7:**
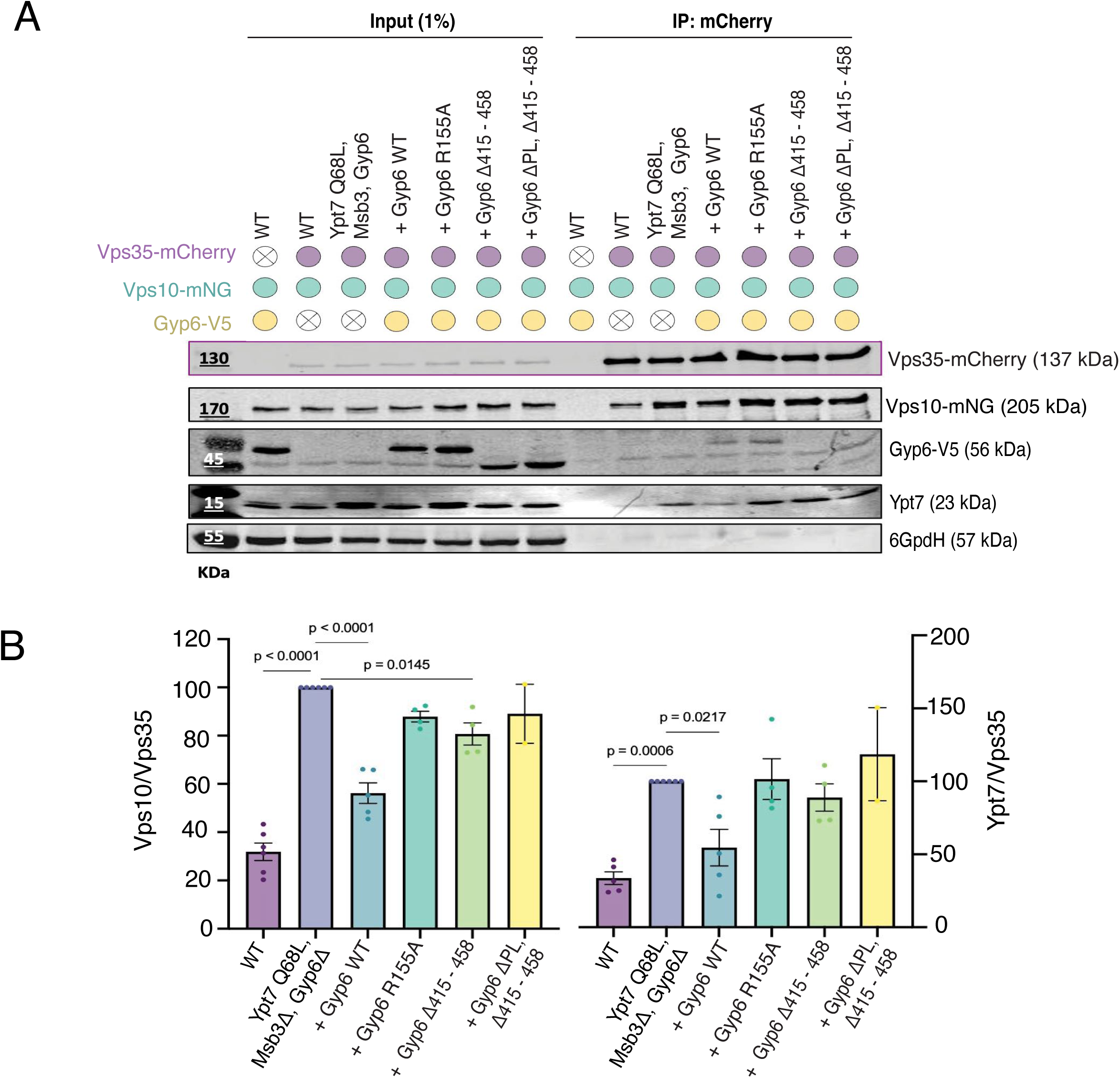
Vps10 cargo and Ypt7 show an increased Retromer association in Gyp6 GAP mutants. **(A)** Co-immunoprecipitation was performed from yeast cell lysates expressing Vps35^yomCherry^, Vps10^mNG^ and Gyp6^V5^. Immunoprecipitates were collected using anti-mCherry beads and probed by immunoblotting with anti-mNG, anti-V5, anti-Ypt7 and anti-mCherry antibodies. Significant increase of both cargo and Rab GTPase with Retromer in Gyp6 mutants. Representative results from at least three independent experiments are shown. Input fractions are shown to assess expression and specificity. (**B**) The amount of Vps10 and Ypt7 co-immunoprecipitated was quantified and normalized to the level of Vps35^mCherry^. Data represent mean ± s.e.m. from *n* = 3 independent experiments. Statistical significance was determined using one-way ANOVA.

**Figure 8:**
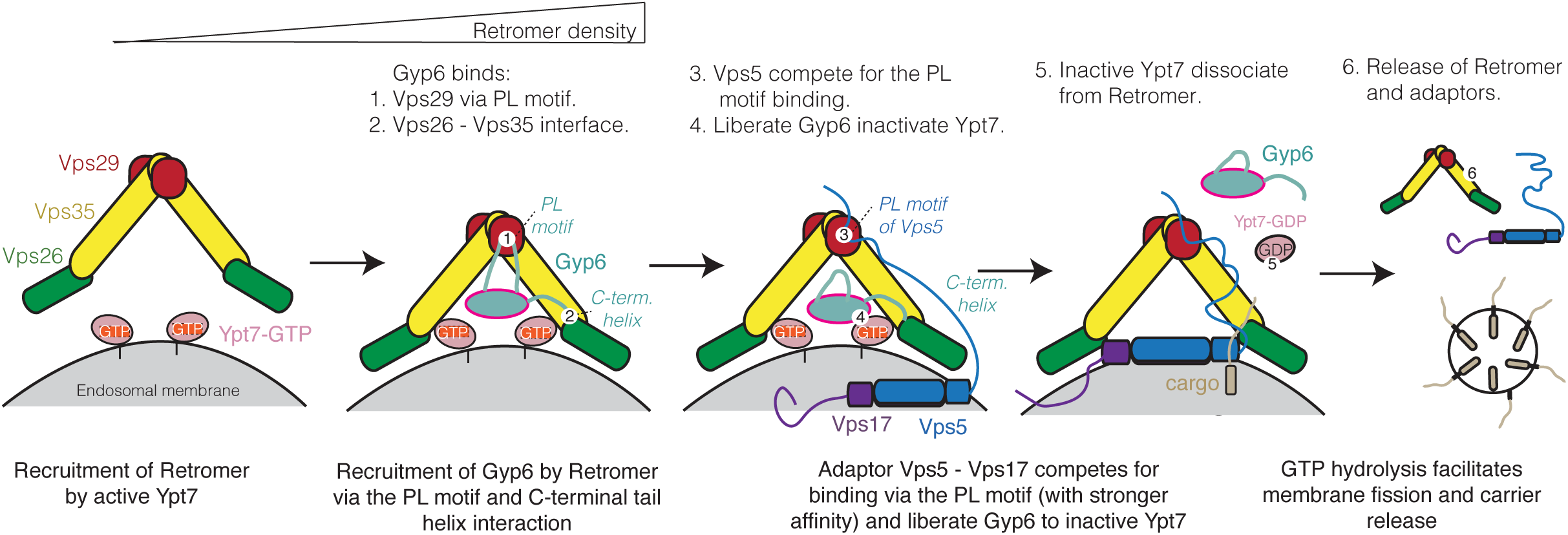

In summary, this suggests that cargo collection and carrier formation by the coat can proceed with active, GTP-loaded Ypt7, but that GTP hydrolysis might be necessary to detach the carrier from the donor membrane. Ypt7 inactivation by GAP activity might then switch Retromer sorting domains from cargo collection to the departure of a carrier.

## Discussion

RAB7 and its yeast homolog Ypt7 function in multiple trafficking pathways and interact with distinct sets of effectors, including tethering factors like the HOPS complex, cargo-sorting complexes such as Retromer, vacuole-mitochondria contact sites, and machinery involved in fusion of vacuoles with endosomes, autophagosomes, or with themselves (Seals *et al*, 2000; Liu *et al*, 2012; Wichmann *et al*, 1992; Hönscher *et al*, 2014; Cabrera *et al*, 2009; Elbaz-Alon *et al*, 2014). Given this functional diversity, we presume that RAB7 and Ypt7 are regulated by multiple GAPs that act in a context-dependent manner to ensure spatial and temporal precision of their inactivation, tailored to the respective function. Our findings provide new insights into the coordination of Ypt7 inactivation and Retromer-mediated cargo sorting, highlighting a role for the GAP protein Gyp6 in coupling GTP hydrolysis with carrier formation. While Retromer has been extensively studied in the context of cargo recognition and membrane remodeling (Harrison *et al*, 2014; Courtellemont *et al*, 2022; Kovtun *et al*, 2018; Cui *et al*, 2018), its regulation by Rab GAPs is relatively poorly understood. Here, we propose that yeast Gyp6 functions as a spatiotemporal regulator of Retromer activity by acting as a Ypt7 GAP during cargo recycling.

GAPs can be promiscuous *in vitro* and, perhaps for this reason, show redundancy in the *in vivo* situation (Lamber *et al*, 2019). This renders the assignment of a GAP to a specific RAB GTPase challenging. Our interaction analyses support an interaction of Gyp6 with Ypt7 in the context of endosomal protein sorting, where both proteins are brought into proximity through their common interaction with Retromer. In line with this, *gyp6*Δ mutants had the strongest impact on endosomal protein sorting by Retromer. Gyp7, a well-characterized Ypt7 GAP that regulates fusion at endosomes and vacuoles (Brett *et al*, 2008; Vollmer *et al*, 1999; Albert *et al*, 1999; Füllbrunn *et al*, 2024), showed no impact on Retromer-dependent Vps10 sorting and only weak signals in the Retromer interaction screen. Gyp3 has a significant effect on Retromer cargo sorting, although its TBC domain showed much weaker GAP activity with Ypt7 than that of Gyp6, in agreement with previous results (Lachmann *et al*, 2012; Albert & Gallwitz, 1999). Since Gyp3 has a well-described role as a GAP for the Rab5 orthologue Vps21 (Lachmann *et al*, 2012; Nickerson *et al*, 2012), it may impact Retromer-dependent sorting indirectly through its influence on endosomal RAB-GTPase conversion. For example, Gyp3 deletion could result in hyperactivation of Vps21 and, consequently, enhanced recruitment of the Ypt7-GEF (Mon1– Ccz1 complex), Ypt7 hyperactivation, activation of endosome-vacuole fusion and a resulting missorting of Vps10 to endosomes (Lachmann *et al*, 2012). This could also explain the additive effect of *gyp3*Δ and ypt7^Q68L^ on Retromer-dependent sorting. While our results do not exclude a function of Gyp3 on Retromer-associated Ypt7, our structural analysis and the presence of dedicated Retromer binding motifs lead us to propose Gyp6 as the genuine GAP of Ypt7 in the context of Retromer.

Our data support the view that Ypt7*^Q68L^*, which is generally presumed to represent a constitutively active form, remains partly functional *in vivo* (Vollmer *et al*, 1999) and that its GTPase activity can be substantially stimulated by GAP proteins to inactivate its effector function on Retromer, albeit at a slower rate. Combining the *ypt7^Q68L^* allele with *gyp3*Δ and *gyp6*Δ leads to the accumulation of Vps10 cargo in clusters at or near the vacuolar membrane that are enriched for both Retromer and Ypt7, and to the accumulation of Retromer complexes with both cargo and Ypt7. Impairing Ypt7 GTP hydrolysis in this way also leads to the accumulation of extended tubules that carry cargo and Retromer, suggesting that while tubule initiation can occur, carrier detachment may require RAB inactivation. This observation supports a model in which Ypt7-GTP promotes cargo clustering and carrier assembly, but its timely inactivation by Gyp6 is necessary for membrane remodeling and carrier release.

While Ypt7 can directly interact with Retromer, the Vps5 – Vps17 SNX-BARs can in turn displace Ypt7 from Retromer (Purushothaman *et al*, 2017; Liu *et al*, 2012; Balderhaar *et al*, 2010). Our previous study also demonstrated that Retromer forms a tight pentameric complex with Vps5 and Vps17 on the membrane via a PL motif in Vps5, which also binds to Vps29 (Chen *et al*, 2025). The Gyp6 PL motif has a lower binding affinity for Vps29 compared to that of Vps5. A stable Gyp6-Retromer complex is hence formed only when the PL motif and the C-terminal tail of Gyp6 are engaged simultaneously. In this conformation, the arginine finger essential for Rab GTP hydrolysis is positioned adjacent to the PL motif – Vps29 interface, far from Ypt7, which is predicted to reside near the N-terminal region of Vps35 (Harrison *et al*, 2014; Gonzalez-Lozano *et al*, 2025). Thus, Gyp6-mediated inactivation of Ypt7 requires a conformational change or dissociation from Retromer. We speculate that Vps5, which binds Retromer with a high affinity, could outcompete the Gyp6 interaction. In this scenario, Gyp6 may remain weakly bound to Retromer through its C-terminal tail, forming a more flexible complex. While Gyp6 remains bound to Retromer, it may inactivate nearby Ypt7 and thereby favor Ypt7 dissociation from Retromer and the formation of the competing Retromer-SNX-BAR complex. This hypothesis is supported by our data, which show that additional deletion of the C-terminal helix of Gyp6 causes a more pronounced defect in Vps10 sorting than deletion of the PL motif alone. A complete sorting defect is observed only when both the PL motif and the C-terminal binding helix are deleted, highlighting the importance of both regions. Given that Gyp6 could be displaced by Vps5 at the PL motif site, we envision a handover mechanism to regulate GAP activity through the assembly of the pentameric Retromer-SNX-BAR complex. Combining our data with the published observations mentioned above, we propose the following model: (1) Active Ypt7 recruits the Retromer to endosomes; (2) With increasing Retromer density, Gyp6 is recruited; (3) Retromer binds cargo such as Vps10 in conjunction with adaptors such as Vps5-Vps17 sorting nexins that bind Retromer through their PL motifs (4); these competing PL motifs liberate Gyp6 to inactivate Ypt7; (6) GTP hydrolysis facilitates membrane fission and carrier release. The catalytic role of Gyp6 and its ability to interact with Retromer thus suggest a mechanism for a Rab-GTPase-controlled switch from coat assembly and cargo collection to carrier release. Given the similarity between Gyp6 and the metazoan GAP TBC1D5 (Jia *et al*, 2016) in their mode of interacting with Vps29 through their PL motifs we assume that this spatiotemporal regulation of carrier formation through GAP handover and Rab-GTPase switching is a general feature of Retromer.

## Materials and Methods

### Yeast Strains and Genetic Manipulations

**(A)** *S. cerevisiae* strains used in this study are listed in **Table S3**. Genetic manipulations were made using the CRISPR-Cas9 approach and confirmed by PCR and sequencing. The plasmids used to carry out CRISPR-Cas9 genome editing were obtained by cloning hybridized oligonucleotides coding for the sgRNA between the XbaI and AatII restriction sites of the parent plasmid (pSP473, pSP474, pSP475, or pSP476, **Table S4**). The oligonucleotides used for hybridization and to control strains by PCR or sequencing are listed in **Table S5**. The integrative plasmids used in this study are listed in **Table S4**. Cells were cultured for transformation at 30°C to mid-log phase in YPD medium (1% [wt/vol] yeast extract, 2% [wt/vol] Bacto Peptone, and 2% [vol/ vol] glucose) or with YNB medium (0.17% [wt/vol] yeast nitrogen base without amino acids and ammonium sulfate, 0.5% [wt/vol] ammonium sulfate, and 2% [vol/vol] glucose) supplemented with the necessary amino acids.

### Proximity labelling

#### Purification of biotinylated proteins via streptavidin beads

Method adapted from (Cho *et al*, 2020). Cells expressing TurboID-V5-Ypt7, *ypt7^Q68L^* or *ypt7^T22N^* (in a BJ3505 *ypt7*Δ background) and TurboID-V5 (in BJ3505 wild-type background) were grown overnight in SC-URA to an OD_600_ of 0.3 (∼ 0.6 × 10^7^ cells). From a freshly prepared biotin stock (100 mM) in dimethyl sulfoxide (DMSO), 100 μM was added to cultures, and cells were grown for another 3 h to a final OD_600_ of 0.8–1. Around 350-400 OD_600_ equivalent units of cells were harvested by centrifugation at 3000 rpm for 10 min, washed twice with ddH_2_O, and cell pellet resuspended in RIPA lysis buffer (50 mM Tris, 150 mM NaCl, 0.1% SDS, 0.5% sodium deoxycholate, 1% Triton X-100, 1x PIC, 1 mM PMSF). Whole-cell lysates were obtained using French Press in RIPA lysis buffer. Free biotin was depleted from the cell extract using Zeba Spin Desalting Columns 7K (Thermo Scientific™ Zeba™ Spin Desalting Columns). Clarified lysates containing around 2.5 mg protein per sample were incubated overnight in 100 µL of pre-equilibrated streptavidin magnetic beads with RIPA buffer (Pierce™ Streptavidin Magnetic Beads, Thermofisher) at 4^∘^C with rotation.

Magnetic beads were subsequently washed in the following order: RIPA buffer (2 min @RT, twice), 1 M KCl (2 min @RT), 0.1 M Na_2_CO_3_ (10 sec @RT), 2 M Urea in 10 mM Tris-HCl pH 8.0 (10 sec @RT) and RIPA buffer (2 min @RT, twice). The beads were then resuspended in fresh RIPA lysis buffer, transferred to a new Eppendorf tube, the buffer was removed, and the dry beads were sent to the Protein Analysis Facility (UNIL) on dry ice for further processing and preparation for LC-MS/MS analysis. For control purposes, 5% of beads were eluted with 4x-Nupage (Thermofisher) supplemented with 2 mM Biotin and 20 mM DTT and boiled for 10 min at 95^∘^C (Vf= 30 µL). 10 µL of eluted sample was loaded into 10% SDS-PAGE gel followed by overnight WB transfer. The Streptavidin-IRDye 680 was used to visualize biotinylated proteins.

### Proteomics analyses

All raw MS data together with raw output tables are available via the Proteomexchange data repository (www.proteomexchange.org) with the accession PXD073331.

#### Protein digestion

Samples were digested following a modified version of the iST method (Kulak *et al*, 2014) (named miST method). 25 μL of miST lysis buffer (1% Sodium deoxycholate, 100mM Tris pH 8.6, 10 mM DTT), were added to the beads. After mixing and dilution 1:1 (v:v) with H_2_O, samples were heated 10 min at 75°C. Turbo-ID beads were first digested with 0.5 μg of trypsin/LysC mix (Promega #V5073) for 1h at 25°C, and sample supernatants were transferred to new tubes. Beads were washed with 50 μL of miST buffer diluted 1/1 in H_2_O, and supernatants were pooled with the previous ones. Reduced disulfides were alkylated by adding 25 μL of 160 mM chloroacetamide (32 mM final) and incubating for 45 min at 25°C in the dark. Samples were then digested overnight at 25°C with 1.0 μg trypsin/LysC mix.

To remove sodium deoxycholate after digestion, two sample volumes of isopropanol containing 1% TFA were added to the digests, and the samples were desalted on a strong cation exchange (SCX) plate (Oasis MCX; Waters Corp., Milford, MA) by centrifugation. After washing with isopropanol/1%TFA, peptides were eluted in 200ul of either 80% MeCN, 19% water, 1% (v/v) ammonia and dried by centrifugal evaporation.

#### Liquid Chromatography-Mass Spectrometry analyses

Data-dependent LC-MS/MS analyses of samples were carried out on a Fusion Tribrid Orbitrap mass spectrometer (Thermo Fisher Scientific) interfaced through a nano-electrospray ion source to an Ultimate 3000 RSLCnano HPLC system (Dionex), via a FAIMS interface. Peptides were separated on a reversed-phase custom packed 45 cm C18 column (75 μm ID, 100Å, Reprosil Pur 1.9 μm particles, Dr. Maisch, Germany) with a 4-90% acetonitrile gradient in 0.1% formic acid at a flow rate of 250 nl/min (total time 140 min). The MS acquisition method used cycled through three compensation voltages (−40, −50, −60V) to acquire full MS survey scans at 120’000 resolution. A data-dependent acquisition method controlled by Xcalibur software (Thermo Fisher Scientific) was set up that optimized the number of precursors selected (“top speed”) of charge 2+ to 5+ from each survey scan, while maintaining a fixed scan cycle of 1.0 s per FAIMS CV. Peptides were fragmented by higher energy collision dissociation (HCD) with a normalized energy of 32%. The precursor isolation window used was 1.6 Th, and the MS2 scans were done in the ion trap. The *m/z* of fragmented precursors was then dynamically excluded from selection during 60 s.

#### Data processing

Data files were analysed with MaxQuant 2.4.7 (Cox & Mann, 2008), incorporating the Andromeda search engine (Cox *et al*, 2011). Cysteine carbamidomethylation was selected as a fixed modification while methionine oxidation and protein N-terminal acetylation were specified as variable modifications. The sequence databases used for searching were the yeast (*Saccharomyces cerevisiae*) reference proteome based on the UniProt database (www.uniprot.org, version of March 1^st^, 2023, containing 6’060 sequences), and a “contaminant” database containing the most usual environmental contaminants and enzymes used for digestion (keratins, trypsin, etc). Mass tolerance was 4.5 ppm on precursors (after recalibration) and 0.5 Da on MS/MS fragments. Both peptide and protein identifications were filtered at 1% FDR relative to hits against a decoy database built by reversing protein sequences.

#### Data analysis

All subsequent analyses were done with an in-house developed software tool (available on https://github.com/UNIL-PAF/taram-backend). Contaminant proteins were removed, and intensity iBAQ values (Schwanhausser *et al*, 2011) were log2-transformed. After assignment to groups, only proteins quantified in at least 4 samples of one group were kept. Missing values were imputed based on a normal distribution with a width of 0.3 standard deviations (SD), down-shifted by 1.8 SD relative to the median. Student’s T-tests were carried out among conditions, with Benjamini-Hochberg correction for multiple testing (adjusted p-value threshold <0.05). Imputed values were later removed. The difference of means obtained from the t-tests were used for 1D enrichment analysis on associated GO/KEGG annotations as described (Cox & Mann, 2012). The enrichment analysis was also FDR-filtered (Benjamini-Hochberg, adjusted p-value <0.02). The -log10 of the adjusted p-value was calculated, and the graphs were generated using GraphPad PRISM software 10.4.0.

### FM 4-64 staining and confocal microscopy

Cells were inoculated from a pre-culture in stationary phase and grown overnight to early mid-logarithmic phase (OD_600_ 0.5 - 0.8) at 30°C, 200 r.p.m. After dilution to an OD_600_ of 0.2 - 0.3 in 1 mL of fresh medium, FM 4-64 (N-(3-triethylammoniumpropyl)-4-(6-(4-(diethylamino) phenyl) hexatrienyl) pyridinium dibromide) in DMSO was added to a final concentration of 10 μM. Cells were stained for 60 min, followed by two washing steps in medium without dye (centrifugation at 3,000 rpm for 2 min) and a subsequent chase of 90 - 120 min (depending on the endocytic capacity of the strain) in 1 mL of fresh medium.

Imaging was performed with a NIKON Ti2 spinning disc confocal microscope with a 100×1.49 NA lens. Imaging was done at room temperature in YNB medium using GFP and mCherry channels with different exposure times according to the fluorescence intensity of each protein. Z-stacks were obtained with a spacing of 0.2 μm between each single focal plane and assembled, when required, into average projections containing five single slices around the middle of the cell. All experiments were repeated at least three times.

To investigate the function of the Retromer, the localization of vacuolar Vps10^mNG^ in different strains was assessed from at least 150 cells per experiment across three independent biological replicates. Mean, SEM calculations from stacks containing around 150 - 200 cells per strain were done in Excel and plots were obtained with GraphPad PRISM software.

### Total cell extracts by TCA

To prepare cell lysates, cells were grown overnight to mid-log phase (OD_600_ 0.5 - 1) at 30 °C from a saturated pre-culture. 2 OD units of cells were harvested by centrifugation (3000 rpm, 3 min). The obtained cell pellet was resuspended in 200 μL of 20% trichloroacetic acid (TCA) and lysed by bead beating with glass beads (0,25 - 0,5 mm, Carl Roth) for 15 min at 4°C using a VIBRAX (IKA). After lysis, 400 μL of 40% TCA was added, supernatant collected and centrifuged at 14,000 rpm for 10 min at RT. Cell lysates were solubilized in 4x-Nupage (containing 1 M Tris and 5% β-mercaptoethanol) at 98°C for 10 min and centrifuged (14,000 rpm, 2 min, RT) to remove cell debris. Equal amounts (0.5 - 0.75 OD_600_ equivalents of cells per sample) were analyzed by Western blot using anti-Ypt7 (homemade), anti- mNeonGreen (32f6-100, Chromotek), anti-mCherry (ab125096, Abcam), anti-V5 (460705, Novex), anti-G6PDH (A9521-1VL, Sigma), anti-Pgk1 (ab113687, Abcam). Detection was carried out using IRDye 680 or 800RD-conjugated secondary antibodies.

### CPY sorting assay

Yeast strains were cultured overnight to achieve saturation in the appropriate medium at 30°C. A 2 μL aliquot of each saturated culture was applied onto YPD agar plates and incubated at 30°C for 16 hours. Following this, the colonies were covered with nitrocellulose membranes and incubated overnight at 30°C. The membranes were subsequently rinsed with double-distilled water (ddH_2_O) and incubated for 1 hour in TBST buffer containing 5% non-fat milk to block nonspecific binding. The membrane was probed for the secreted protein carboxypeptidase Y (CPY) using an anti-CPY mouse monoclonal antibody (ab113685; Abcam) for 4 hours at room temperature. Detection was carried out using an IRDye 800RD-conjugated goat anti-mouse secondary antibody (926-32210; LI-COR).

### Immunoprecipitation

The interaction between cargo (Vps10), Retromer, and Ypt7 was analyzed following a protocol adapted from the study (Suzuki *et al*, 2019). Briefly, cells expressing Vps10^mNG^ and Vps35^yomCherry^, from their endogenous promoter, were cultured overnight until they reached the mid-logarithmic phase (OD_600_ between 0.8 and 1.0), starting from a saturated pre-culture.

A total of 50 mL of cells were collected, washed twice with immunoprecipitation (IP) buffer (20 mM Hepes-KOH, pH 7.2, 0.2 M sorbitol, 50 mM AcOK, 1 x protease inhibitor cocktail, 1 mM PMSF), and then harvested. The cells were lysed with 0.5-mm zirconia beads in 250 µL of IP buffer for 5 minutes at 4°C using a VIBRAX (IKA). To this lysate, 250 µL of IP buffer containing 2.0% saponin was added, to obtain a final concentration of 1.0%. The samples were then rotated at 4°C for 60 minutes at 10 rpm. The solubilized lysates were clarified by centrifugation at 500 g for 5 minutes at 4°C, followed by high-speed centrifugation at 17,500 g for 10 minutes. The resulting supernatants were normalized to 10 mg of total protein per sample, measured by Nanodrop, and incubated with pre-equilibrated anti-RFP magnetic agarose beads (RFP-Trap Magnetic Agarose from Chromotek) at 4°C for 3 h. After incubation, the beads were washed three times with IP buffer containing 1.0 % saponin. Before the final washing step with IP buffer, beads were transferred into a new Eppendorf tube. The bound proteins were eluted by heating the beads in 4 x Nupage buffer (containing 5% β-mercaptoethanol, 10 mM DTT, 1 mM PMSF, and 1 x protease inhibitor cocktail) at 95°C for 10 minutes. Samples were loaded into a freshly prepared SDS-PAGE gradient gel (4-18%).

To evaluate the interaction between Retromer and Gyp6, a co-immunoprecipitation protocol adapted from the previously described method was employed with minor modifications. Yeast cells expressing Gyp6^V5^ (or its variants) and Vps35^yomCherry^, under endogenous promoter, were cultured overnight from saturated pre-cultures and harvested at mid-log phase (OD₆₀₀ = 0.8– 1.0). A total of 75 mL of culture was collected, washed twice with TGN buffer (50 mM Tris-HCl, pH 7.4, 100 mM NaCl, 5% [v/v] glycerol, 1x protease inhibitor cocktail, 1 mM PMSF), and pelleted by centrifugation. Cell lysis was performed at 4°C using 0.5-mm zirconia beads in 300 µL of TGN buffer for 5 minutes using a VIBRAX (IKA). Subsequently, 300 µL of TGN buffer containing 1.0% Triton X-100 was added to the lysate to achieve a final detergent concentration of 0.5%. Samples were rotated at 4°C for 60 minutes at 10 rpm to facilitate membrane solubilization.

The lysates were clarified by centrifugation at 500 x g for 5 minutes at 4°C, followed by a second centrifugation step at 17,500 x g for 10 minutes. The resulting supernatants were adjusted to a final protein concentration of 10 mg in a total volume of 500 µL (quantified via NanoDrop) and incubated with pre-equilibrated anti-RFP magnetic beads (RFP-Trap Magnetic Particles, Chromotek) for 2 hours at 4°C. Following incubation, beads were washed three times with TGN buffer containing 0.5% Triton X-100. Before the final wash with detergent-free TGN buffer, the beads were transferred to fresh microcentrifuge tubes. Bound proteins were eluted by incubating the beads in 4× NuPAGE sample buffer (supplemented with 5% β-mercaptoethanol, 10 mM DTT, 2 mM PMSF, and 1 × protease inhibitor cocktail) at 95°C for 10 minutes. Eluted proteins were resolved by SDS-PAGE using freshly cast 4–18% polyacrylamide gradient gels.

### Western blot

All Western blot analysed in this study were performed using a LI-COR Odyssey SA system to detect secondary antibodies conjugated with fluorescent labels. The intensities of the corresponding bands were quantified using FIJI software. All measurements were carried out across a minimum of three independent experiments, as described in the figure legends. All the calculations were done in Excel, and graphs were generated using Prism 10.4.0.

### Molecular biology and cloning for bacteria expression

cDNA encoding full-length ctVps29 (uniport number: G0RZB5), ctVps26 (uniport number: G0S0E6) and ctVps35 (uniport number: G0S709) were cloned into pET28a and pGEX6P1 vectors similar to the one described previously (Chen *et al*, 2025). ctVps29 for the ctGyp6L (no uniport number, GenBank: KAH6850218.1) binding experiment was also cloned into pET28a vector but with an uncleavable C-terminal His6-tag. All the GAP and Ypt7 constructs in this study including ctGyp3^TBC^/ctMsb3^TBC^ (uniport number: G0S722), ctGyp6L^TBC^ (residues 1 to 374), LsmA-ctGyp6L^676-773^, ctGyp7^TBC^ (uniport number: G0SFD4), Gyp6, ctYpt7 (uniport number: G0SGE1), ctVps21 (uniport number: G0SHN0), ctRab5like (uniport number: G0SHY4) and Ypt7 were cloned into pGEX6P1 vector. For ctYpt7, ctVps21, ctRab5like and Ypt7, the C-terminal Cys, Ser and Cys were substituted with His_6_-tag to mimic the prenylation. Site-directed mutagenesis was performed to generate mutant constructs with custom-designed primers. All constructs were verified using DNA sequencing.

### Recombinant protein expression and purification in bacteria

All the bacterial constructs including ctRetromer, ctGyp3^TBC^, ctGyp6L^TBC^, ctGyp7^TBC^, ctYpt7, ctVps21, ctRab5like, LsmA-ctGyp6L^676-773^, Gyp6, Ypt7 and associated mutants were expressed in BL21 (DE3) competent cells using autoinduction method as described previously (Chen *et al*, 2025). To produce ctRetromer trimer complex, GST-tagged ctVps29 and untagged ctVps35 co-expressed cell pellet were mixed with N-terminal His-tagged ctVps26 and resuspended in lysis buffer containing 50 mM Tris-HCl pH 7.5, 200 mM NaCl, 50 μg/ml benzamidine and DNase I before passed through a Constant System TS-Series cell disruptor. To generate ctRetromer specifically for ctGyp6^TBC^ binding experiment, C-terminal His_6_-tagged ctVps29 and GST-tagged ctVps35 co-expressed cell pellet were used. As we found the GST and the PreScisison protease recognition site on the N-terminus of ctVps29 likely affect ctGyp6 from binding. For all constructs, the lysate was clarified by centrifugation and the supernatant was loaded onto either Talon® resin (Clontech) or glutathione Sepharose (GE healthcare) for initial purification. For ctRetromer, both Talon® resin (Clontech) and glutathione Sepharose (GE healthcare) were applied to obtain the correct stoichiometry ratio of the complex. On-column PreScission protease cleavage of GST tag was performed overnight at 4°C using protocol described previously (Chen *et al*, 2025). The flow-through containing the fusion tag cleaved protein was further purified using size-exclusion chromatography (SEC) using either HiLoad® Superdex 200 or Superdex 75 16/600 columns equilibrated with a buffer containing 50 mM Tris-HCl pH 7.5, 200 mM NaCl, 2 mM β-ME.

To obtain the active and inactive form of Ypt7 constructs, the purified wild-type ctYpt7 and Ypt7 in SEC buffer were treated with 1 mM EDTA and 1 mM DTT. After 1 hour incubation on ice, 1.5 mM of either non-hydrolyzable GTP analog GppCp (Abcam) or GDP was added to the sample together with 10 mM MgCl_2_ and incubated on ice for further 1 hour. The reaction was terminated by passing the sample mixture onto an analytical Superdex 75 10/300 increase column using the SEC buffer described above.

### Crystallization and data collection

All crystals were grown by hanging-drop vapor diffusion method at 20°C. Diffraction quality crystal of ctGyp6L^TBC^ was obtained using a protein concentration of 12 mg/ml in condition composed of 0.1 M Bis-Tris pH 6.5, 25% PEG 3350 and 5% glycerol. For the co-crystallization, 3 molar excesses of ctGyp6L^87-107^ peptide or Gyp6^105-122^ peptide was added to the purified mVPS29 at a final concentration of 11 mg/ml. Diffraction crystal of mVPS29 – ctGyp6L^87-107^ peptide was obtained in condition containing 0.2 M sodium fluoride, 18% PEG 3350 and 5% glycerol. For the mVPS29 – Gyp6^105-122^ peptide crystals, the best condition composed of 0.1 M Tris-Base pH 8.5, 22% PEG 3350. Crystals grew after 14 days were directly cryoprotected in reservoir solution except for the Gyp6 peptide containing crystals where an additional 5% glycerol were added. X-ray diffraction data were carried out at the Australian Synchrotron MX2 beamline at 100 K.

### Structure determination

The data collected were indexed and integrated using AutoXDS and scaled using Aimless (Evans & Murshudov, 2013; Kabsch, 2010). Both structures were solved by molecular replacement (MR) using Phaser (Mccoy *et al*, 2007). In the case of ctGyp6L TBC domain crystal, a model generated by Alphafold2 were applied as the template. For the Gyp6 peptide containing crystal structures, the wild-type human VPS29 structure (PDB ID: 6XS7) was used as the template. The initial electron density map obtained after MR guided the locations of the mVPS29, ctGyp6L TBC domain and the peptides. Structure refinement was performed with PHENIX beginning with rigid-body refinement (Adams *et al*, 2010), Translation-Libration-Screw (TLS) and isotropic B factors. The program Coot was used to inspect and fitting of resulting model guided by Fo–Fc difference maps. Ordered water molecules were first automatically included in the model, followed by manual inspection. The coordinates for small ligands such as glycerol, PEG and β-ME were generated using LIBCHECK from Coot. Molprobabity was used to evaluate the quality and geometry of the refined structures (Williams *et al*, 2018). Data collection and refinement statistics are summarized in **Table S2**. Molecular figures were generated using PyMOL and structural comparison was performed using FALI and Foldseek (Kempen *et al*, 2023). Sequence conservation was mapped on the structure with ConSurf server (Ashkenazy *et al*, 2016).

### Isothermal Titration Calorimetry

ITC experiments on ctVps29 and Gyp6 interactions were performed using a Microcal PEAQ ITC (Malvern) at 30°C. All the proteins and peptides including ctVps29, ctGyp6L^87-107^, ctGyp6L^TBC^ and Gyp6^105-122^, used for ITC experiments were prepared in buffer containing 50 mM Tris-HCl pH 7.5, 200 mM NaCl, 2 mM β-ME. To confirm the binding of Gyp6 PL motif is required for binding to Retromer, 350 µM of Gyp6^105-122^ was titrated into 13 µM of ctVps29. In the case of ctGyp6 and ctRetromer binding experiment, 1000 µM of ctGyp6L^TBC^ was titrated into 40 µM of ctVps29. The effect of cyclic peptides on blocking the PL motif-mediated binding was carried out by adding 5 molar excess RT-D3 into ctVps29 prior of the titrating experiments. To investigate the binding between ctGyp6L tail and ctRetromer, 400 µM - 600 µM of ctGyp6L^680-698^ or ctGyp6L^760-773^ peptides were titrated into 14 µM of ctRetromer. The binding experiment using chimera construct was done by titrating 580 µM of either wild-type or IL/F mutant of LsmA-ctGyp6L^676-773^ into 13 µM of ctRetromer. In all cases, the ITC experiments were performed with a single injection of 0.4 μl (not used in data processing) followed by 12 series injections of 3.22 μl each, spaced 180 sec intervals and stirred at 750 rpm. The thermodynamic parameters of binding including the dissociation constants (Kd,), enthalpy (ΔH), and Gibbs free energy (ΔG) were obtained after fitting the integrated and normalized data to a single-site binding model. The stoichiometry (N) was refined initially, and the if the value obtained was close to 1, then, N was set to exactly 1.0 for calculation. Data presented are the mean of triplicate titration for each experiment.

### AlphaFold Predictions

All protein models displayed in the manuscript were generated using either AlphaFold2 Multimer implemented in the Colabfold interface (Mirdita *et al*, 2022) or AlphaFold3 server (https://alphafoldserver.com/) available on the Google platform. The default settings for ColabFold were used to construct multiple sequence alignments using MMseqs2 (Mirdita *et al*, 2019). For all modelling experiments, we assessed (i) the prediction confidence measures (pLDDT and interfacial iPTM scores), (ii) the plots of the predicted alignment errors (PAE) and (iii) backbone alignments of the final structures. The screening of Ypt7 and the TurboID captured interactors was done using batch mode. The confidence level of scRetromer and Gyp proteins screening were calculated using the sum of IPTM score generated by AlphaFold3 and AlphaFold2. For clarity, the confidence level score was differentiated into two groups as full-length scRetromer (as X-Y plot) and individual subunits (as heatmap). All the graphs were generated using Prism 10.4.0.

### GTPase activity assay

GTPase activity was performed in the GTPase-GloTM assay (Promega) using manufacturer’s instructions. In brief, 5 μl of protein/protein mixtures (in general, 1 to 2 μM of Ypts, 0.8 to 1 μM GAPs and 3 μM ctRetromer) was mixed with 5 μL of GTP-solution composed of 10 μM GTP, 1 mM DTT, 50 mM Tris-HCl pH 7.5, 100 mM NaCl, 10 mM MgCl_2_ in 384-well plates. To initiate the GTP reaction, the reaction mixture was incubated for 70 to 80 min at 25°C. The reaction was terminated by adding 10 μL of GTP-Glo buffer, which contains a nucleoside-diphosphate kinase that converts remaining GTP in the solution to ATP. The measurement was carried out by adding 20 μL of detection reagent containing a luciferase/luciferin mixture, making the ATP luminescent. In every set of assay experiments, a GTP-solution buffer sample without protein was used as the control and for normalization (represented as 0% GTP hydrolysis). Reactions were carried out with at least 3 replicates per assay experiment to calculate the average and standard deviation. All the calculation was done in Excel, and graphs were generated using Prism 10.4.0.

### Mass photometry

Molecular mass measurement of LsmA-ctGyp6L^676-773^ was performed using a Refeyn OneMAP mass photometer (Refeyn Ltd) following manufacturer’s instructions. In brief, the calibration was formed first using protein standards with known molecular weight (i.e., 66, 132 and 480 kDa) in buffer containing 50 mM Tris-HCl pH 7.5, 200 mM NaCl, 2 mM β-ME. Next, 3 μL of purified LsmA-ctGyp6L^676-773^ at a final concentration of 50 nM – 100 nM was loaded onto the mass photometer, and 100 frames were recorded. The precise molecular mass was calculated using the calibration standards measured as reference.

## Supporting information

Supplementary Datasheet

## Acknowledgements

We thank the Protein Analysis Facility of the Faculty of Biology and Medicine, University of Lausanne, Lausanne, Switzerland, for mass spectrometry-based protein analytics. We acknowledge the use of the Protein Analysis facility of UNIL, of the University of Queensland Centre for Microscopy and Microanalysis (CMM) Facility, and the assistance of K. A. Arachchige. X-ray data were collected on the MX2 beamline at the Australian Synchrotron. B.M.C. is supported by an Investigator Grant from the National Health and Medical Research Council (APP2016410). AM has been supported by grants from the SNSF (31003A_179306, 310030_204713, and 10.006.083).

## Author contributions

A.C.A., K.-E.C., B.M.C., and A.M. conceived the study. A.C.A., performed cell-based experiments and imaging. Protein preparation was performed by A.C.A., K.-E.C., Q.G, and S.E. K.-E.C., performed biochemical characterization, assays, protein crystallography, and structural analysis. K.-E.C., and M.H., performed in silico screening and modelling. M.L., performed peptide synthesis. A.C.A., K.-E.C., B.M.C., and A.M. supervised the research. Manuscript and figures were prepared by A.C.A., K.-E.C., B.M.C., and A.M.

## Conflicts of Interest

The authors declare that they have no conflict of interest.

## Supplementary Information

**Suplementary Figure 1.**
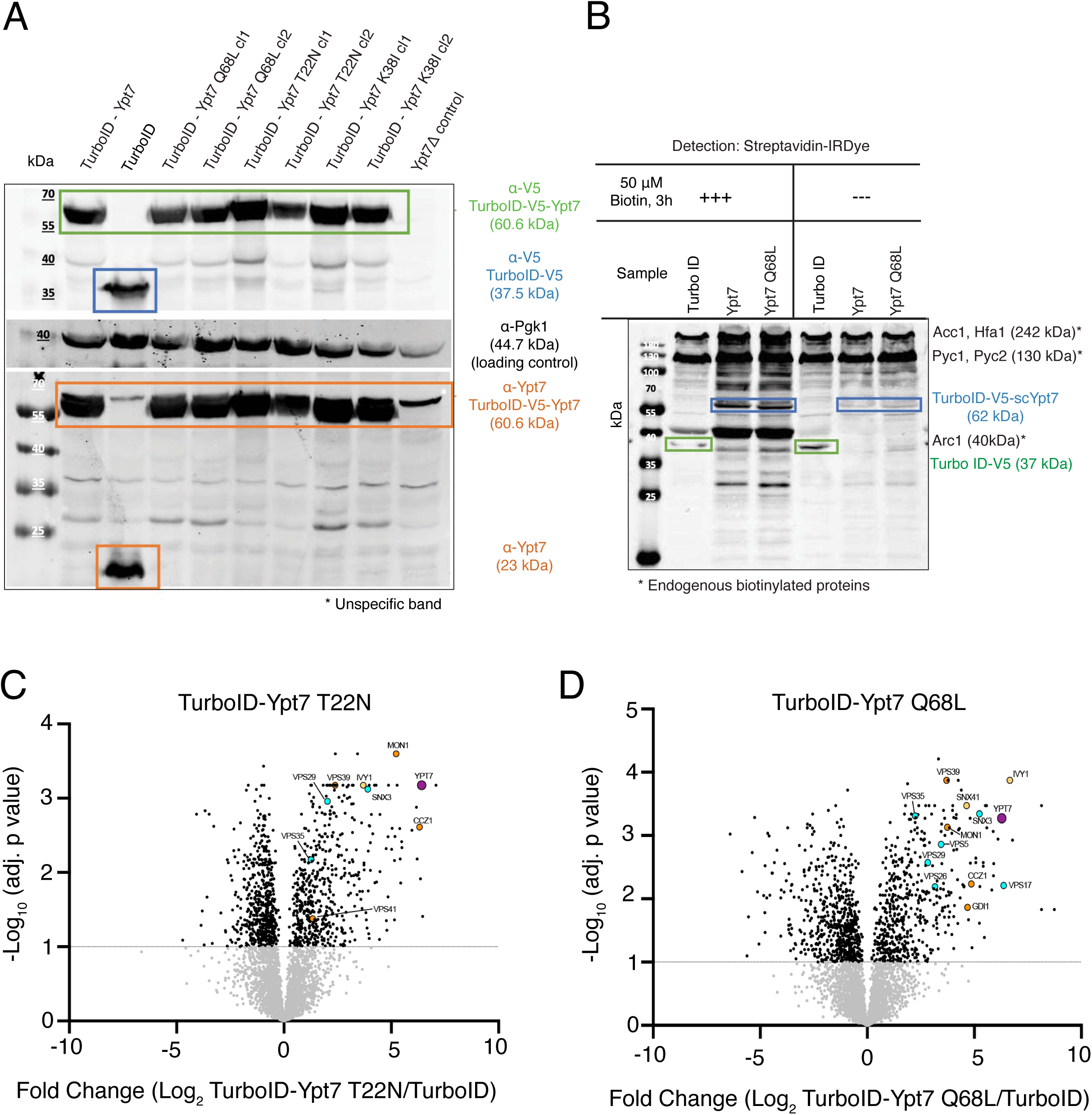
TurboID-V5-Ypt7 protein quantification. (A) Western blot analysis of TurboID-V5-Ypt7 (wt, Q68L and T22N variants) expression in yeast cells. Whole-cell lysates were probed with anti-V5 and anti-Ypt7 antibodies to detect TurboID fusion proteins. Comparable expression levels were observed across samples, ensuring differences in biotinylated interactors reflect proximity and not variation in TurboID expression. PGK1 served as loading control. A representative blot from three independent experiments is shown. (B) Immunoblot of streptavidin precipitates, before and after the addition of biotin, from cells expressing TurboID-V5-Ypt7 and TurboID-V5-ypt7*^Q68L^*. Volcano plots of (C) TurboID-ypt7^T22N^ and (D) of TurboID-ypt7^Q68L^ proximity labelling versus control cells expressing only cytosolic TurboID, from a proteomics analysis of streptavidin-biotin pulldowns.

**Supplementary Figure 2.**
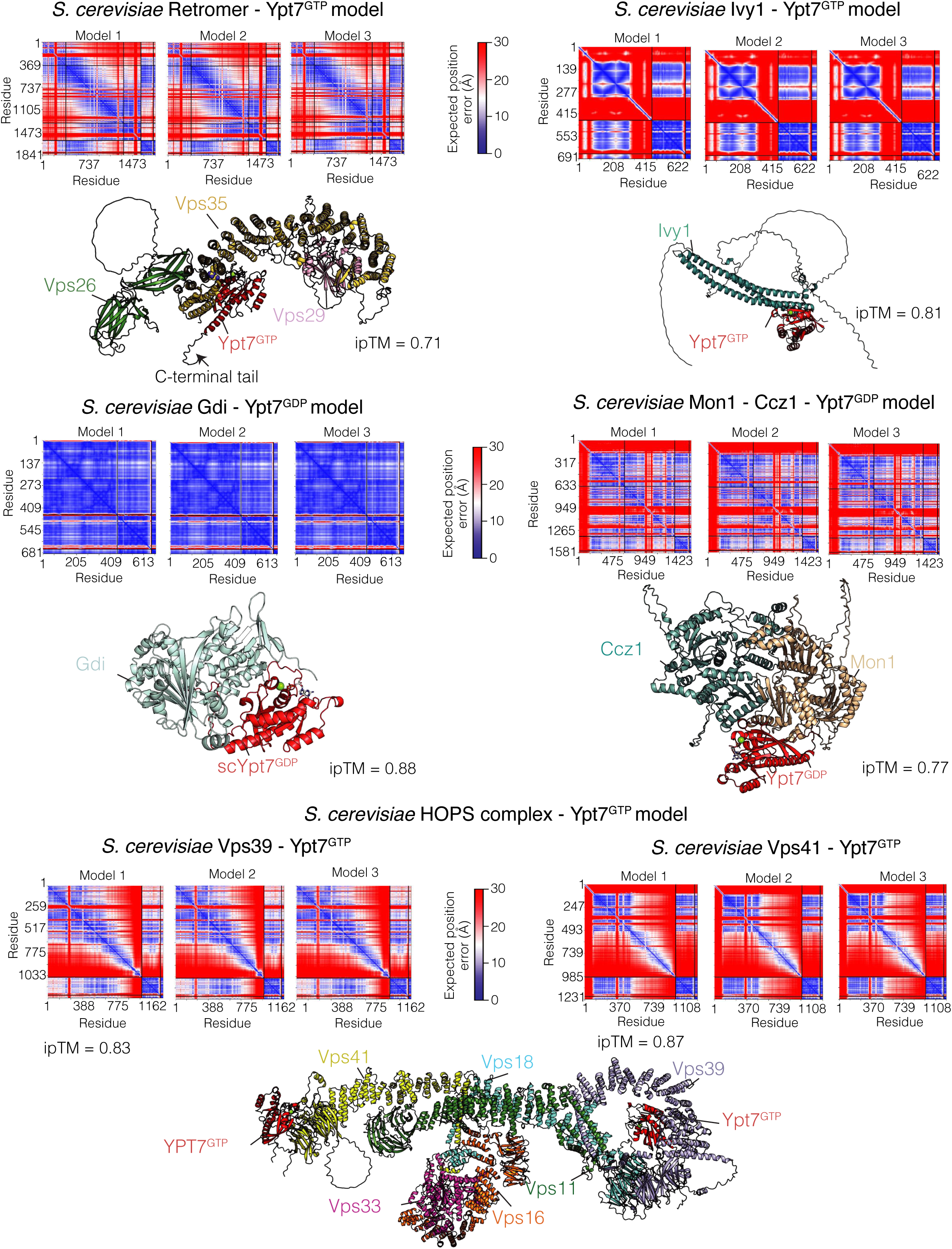
In silico screening identified direct interactors of Ypt7. In all cases. the top three ranked models are aligned and shown in ribbon. The models are coloured according the pLDDT confidence score. Panel on the top of the models shows the predicted alignment error (PAE) plots for the top-ranked prediction of the complex. The ipTM score represent the averaged iPTM score from the top three-ranked models. HOPS complex – Ypt7 model was generated by combining the prediction from multiple subcomplexes including Vps11 – Vps18 – Vps41^729-992^ complex, Vps18^626-918^ – Vps16 – Vps33 – Vps41^729-992^ complex, Vps11^559-1029^ – Vps18^1-658^ – Vps39 – YPT7^GTP^ complex and Vps41 – Ypt7^GTP^ complex. All 4 models were combined using the previous CryoEM structure (PDB ID: 7ZU0) as the template.

**Supplementary Figure 3:**
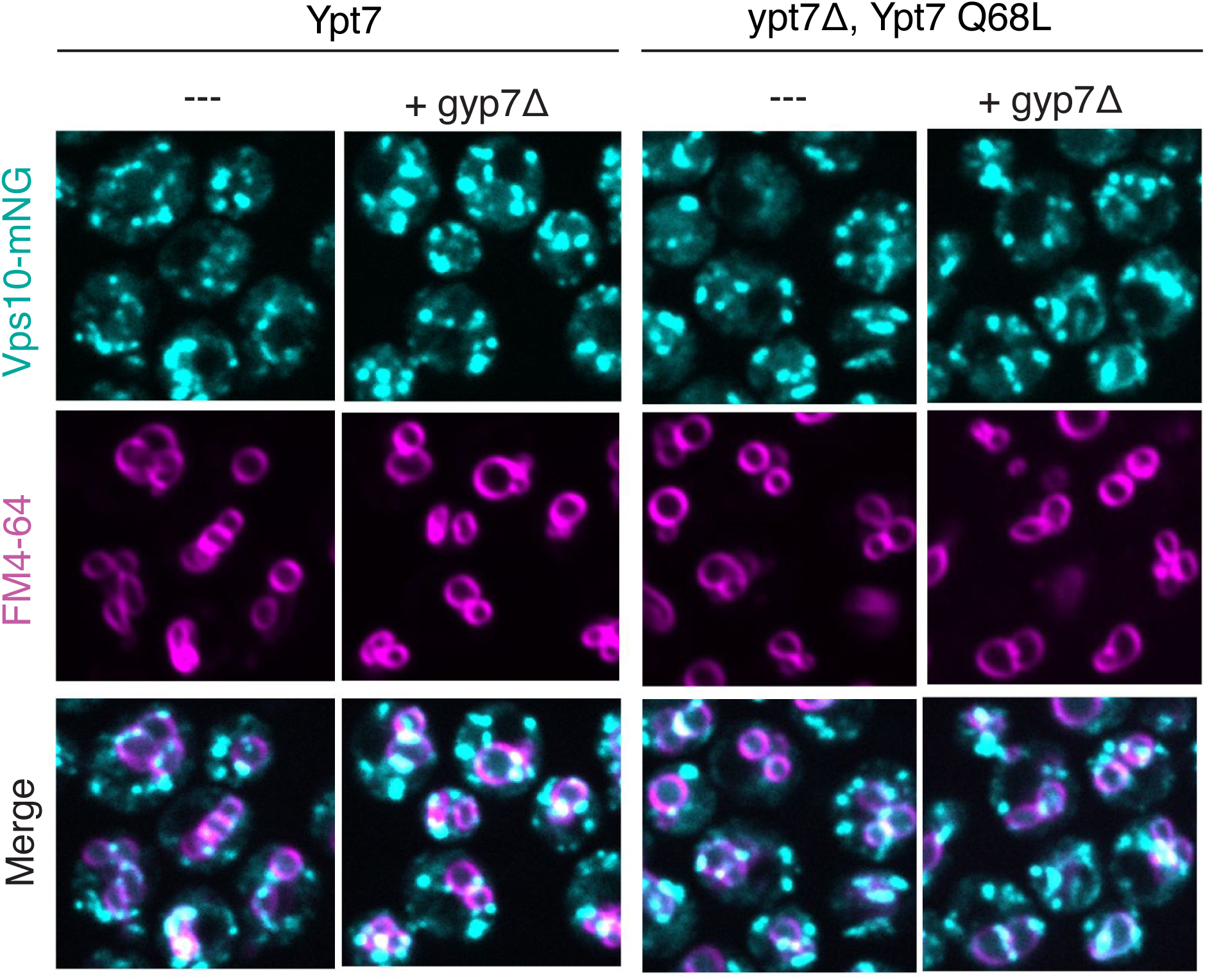
Effect of *gyp7*Δ mutations on Vps10-mNG localisation in the ypt7 Q68L background. Live-cell fluorescence microscopy of yeast expressing Vps10^mNeonGreen^ (cyan) following FM 4-64 pulse-chase labelling to mark the vacuolar membrane (magenta) as shown in Figure 2A.

**Supplementary Figure 4:**
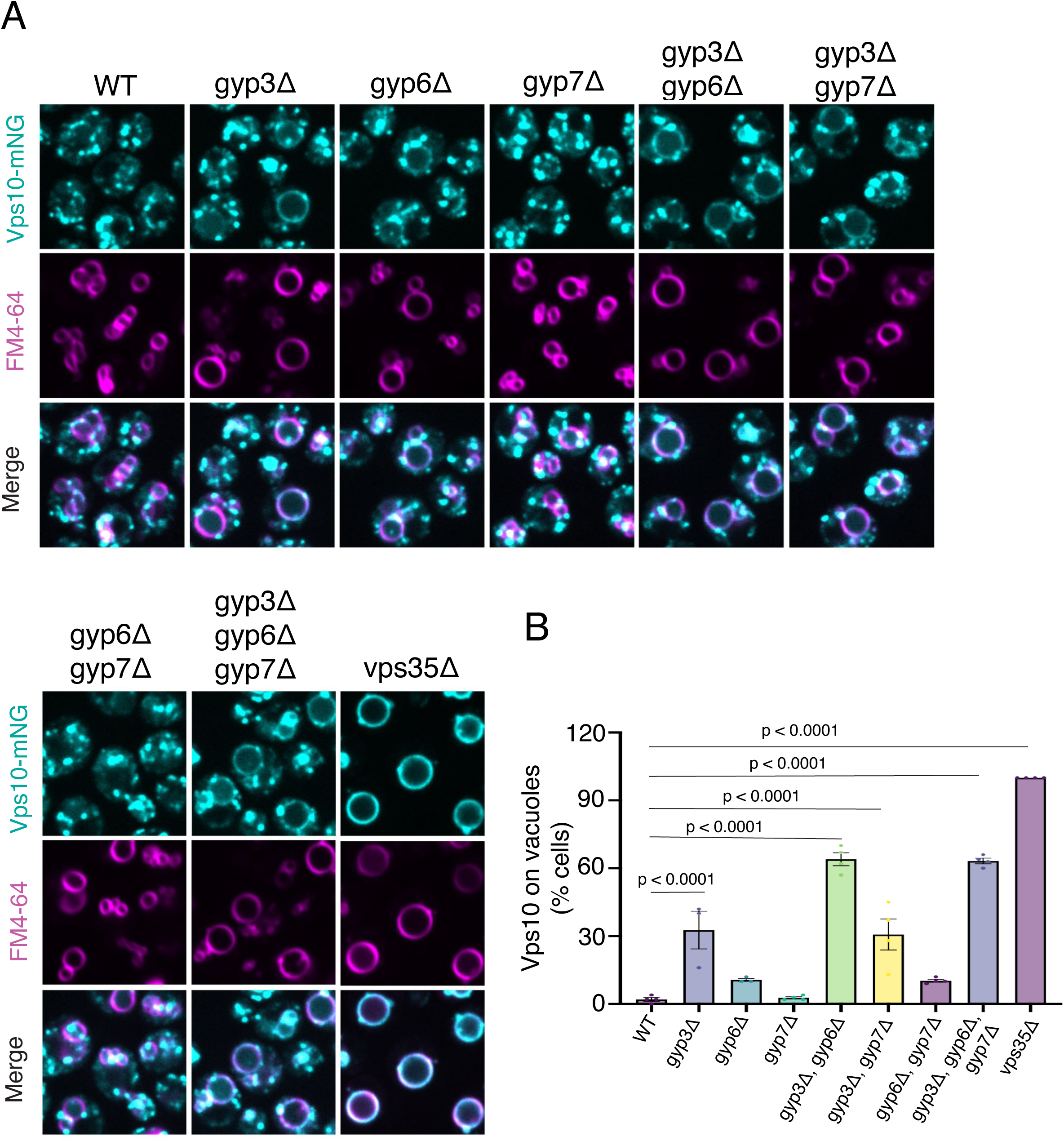
Effect of *gyp7*Δ mutations on Vps10-mNG localisation. **(A)** Cells carrying the indicated combinations of gene deletions and genomic fluorescent tags were cultured, imaged and **(B)** quantified as in Figs. 2A and 2B.

**Supplementary Figure 5.**
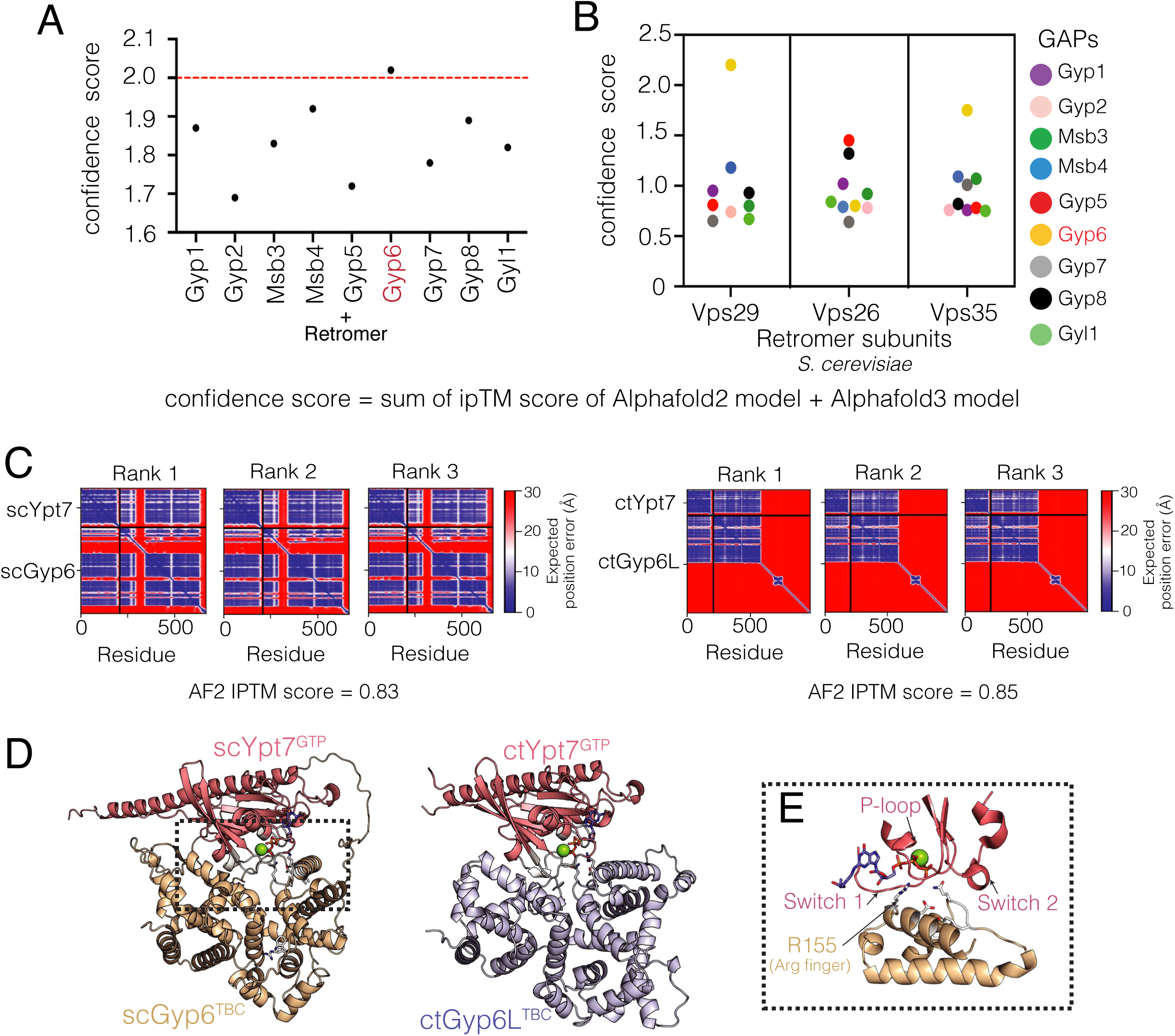
Ypt7 and Retromer are both predicted to bind Gyp6. (A) In silico screening of *S. cerevisiae* Retromer and Rab GAPs. The confidence score of potential interactors was assessed based on the sum of the iPTM score averaged from three AF2 and AF3 models. (B) Among the 9 yeast TBC domain containing GAPs, Gyp6 was confidently predicted to bind to Vps29 subunit of Retromer. (C) The PAE plots for the top ranked models of Ypt7 – Gyp6 and ctYpt7 – ctGyp6L complexes predicted by AlphaFold2. The iPTM score represents the averaged iPTM score from the three models. (D) Cartoon representation of the top ranked Ypt7 – Gyp6 and ctYpt7 – ctGyp6L complexes. GTP and Magnesium in the active site of Ypt7 were modelled based on the AF3 prediction. (E) Close view of Ypt7 – Gyp6 binding interface showing that Gyp6 recognize Ypt7 through canonical interaction.

**Supplementary Figure 6.**
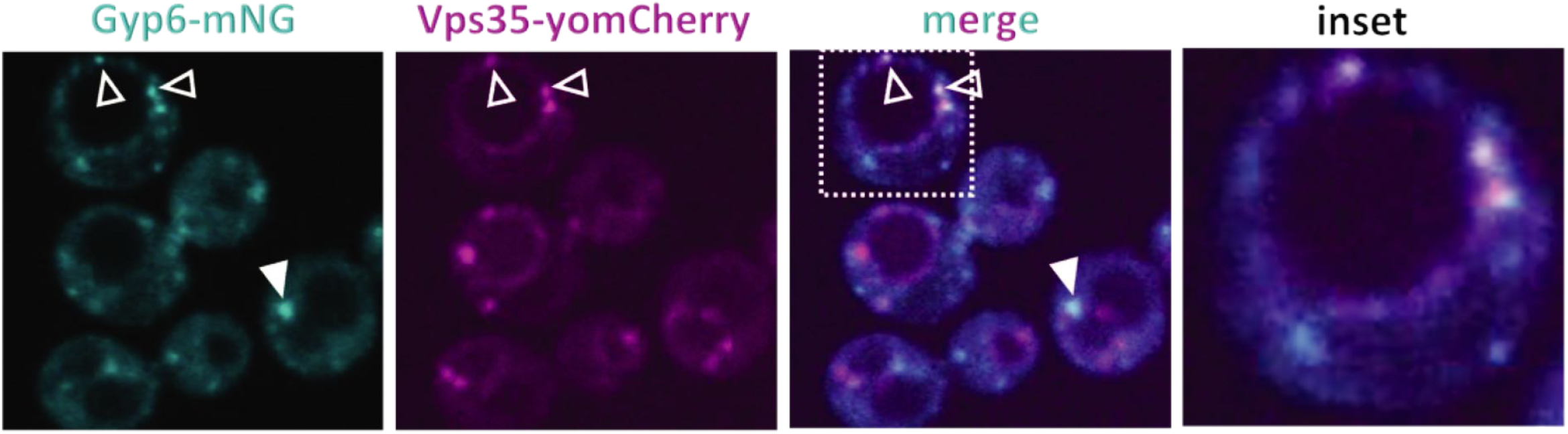
Gyp6 – Retromer colocalization. SEY6210 cells carrying genomically tagged Gyp6^mNG^ and Vps35^yomCherry^ were analysed by spinning disc fluorescence microscopy. Arrowheads indicate Gyp6^mNG^ and Vps35^yomCherry^ colocalising in the same structures. Scale bars: 5 μm.

**Supplementary Figure 7.**
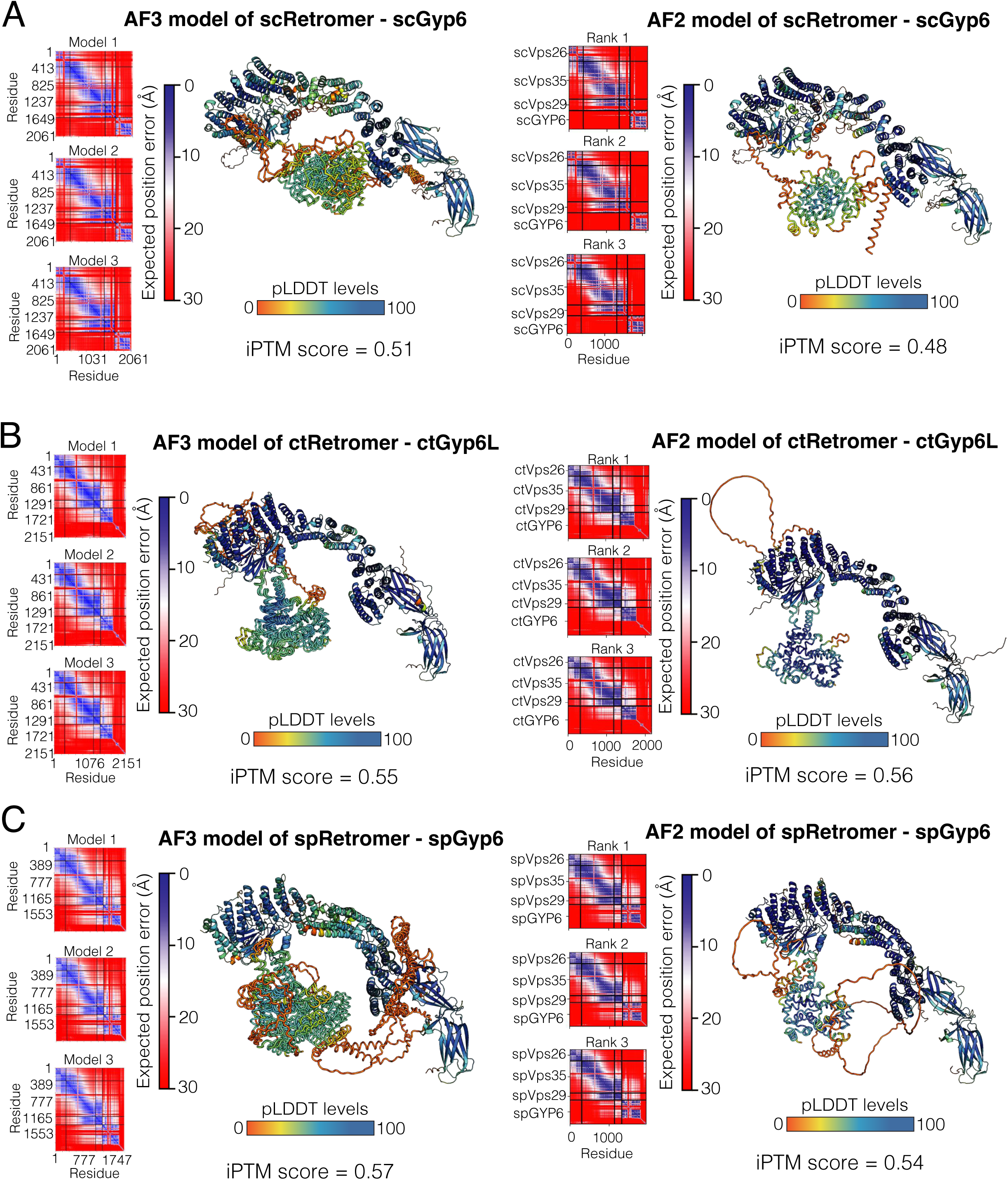
AlphaFold modelling of yeast Retromer and Gyp6 complex. **(A)** AF3-(left) and AF2-(right) predicted complex between Retromer and Gyp6, **(B)** ctRetromer and ctGyp6L, and **(C)** spRetromer – spGyp6. For AF3 prediction, the top three ranked models are aligned and shown in ribbon diagram. For AF2 prediction, only the top ranked model is shown. The models are coloured according the pLDDT confidence score. Panel on the left of the models shows the predicted alignment error (PAE) plots for the top-ranked prediction of the complex. The iPTM score represent the averaged iPTM score from the top three-ranked models.

**Supplementary Figure 8.**
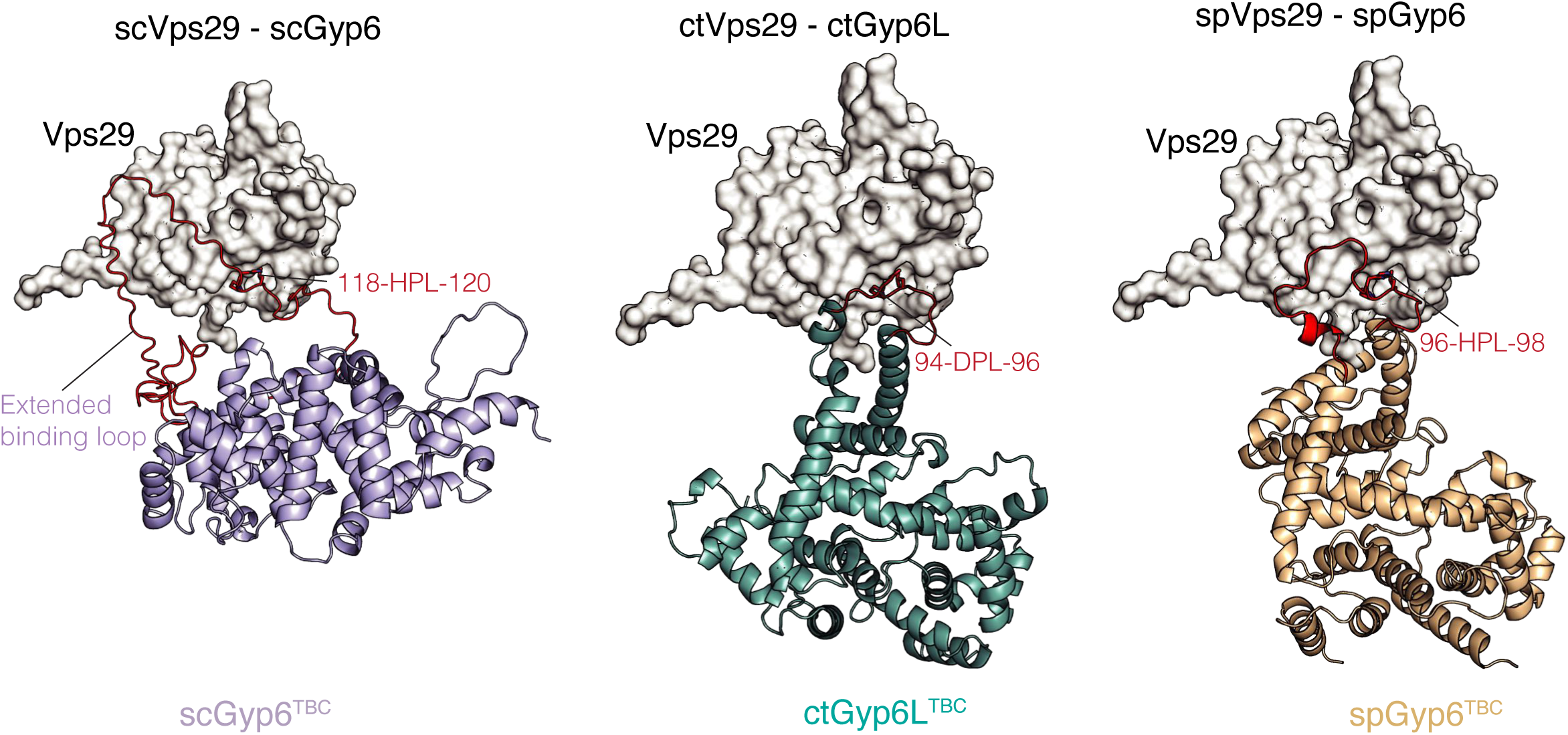
Comparison of the Vps29 - Gyp6^TBC^ models from three different yeast species. While all three Gyp6 models (left: Gyp6, middle: ctGyp6L, right: spGyp6) show a conserved TBC domain fold, they vary in the length of the loop containing the PL motif (highlighted in red).

**Supplementary Figure 9.**
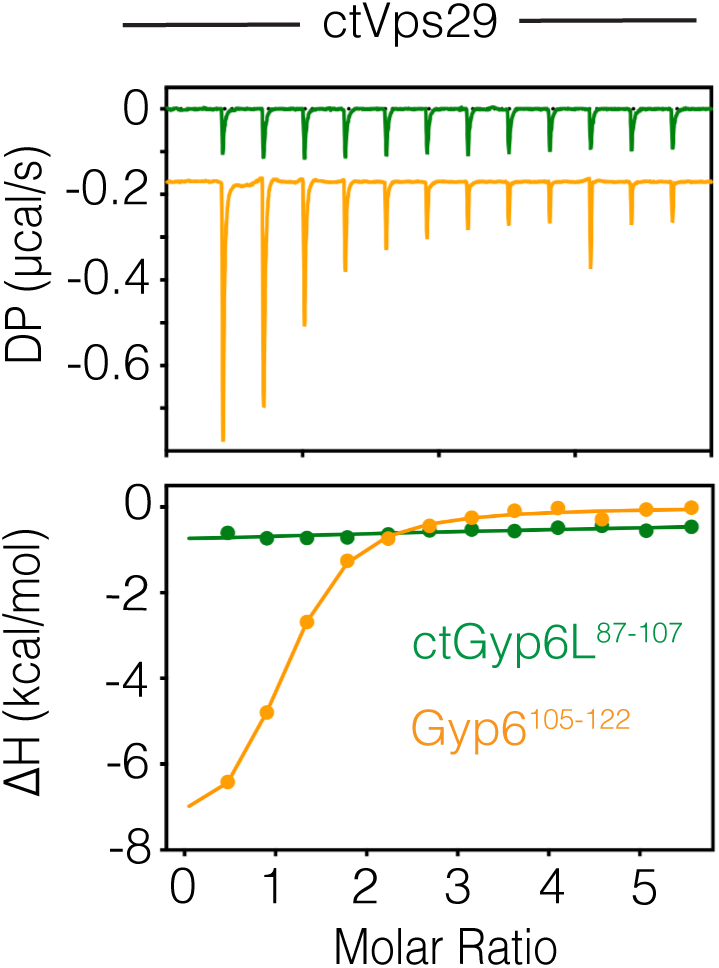
ITC measurement of Gyp6^105-122^ and ctGyp6L^87-107^ peptides binding to ctVps29. ITC graph shows the integrated and normalized data with a 1 to 1 ratio binding ratio.

**Supplementary Figure 10.**
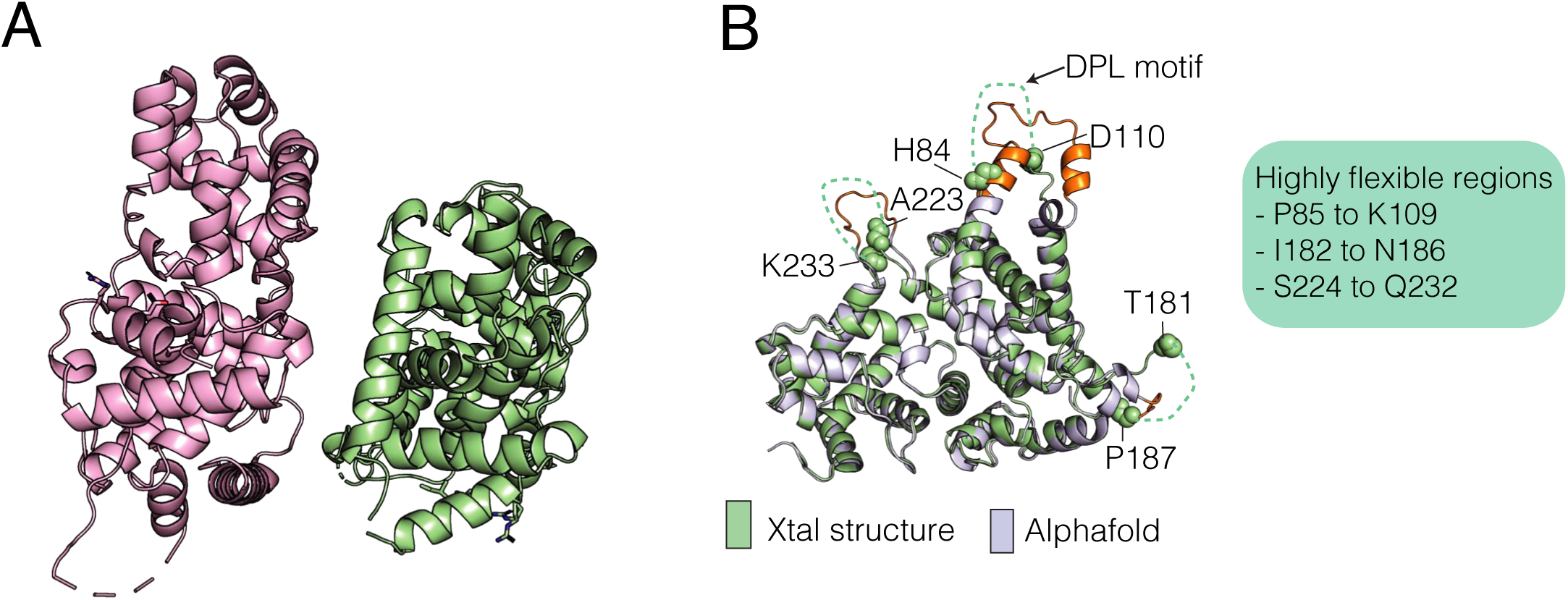
Crystal structure of ctGyp6L^TBC^ shows three distinct disordered loops. (A) Crystal structure of ctGyp6L^TBC^ solved at 2.1 Å resolution reveals two molecules in asymmetric unit. (B) Superimposition of the crystal structure and AF2 model highlighting the three distinct disordered loop observed in the structure.

**Supplementary Figure 11.**
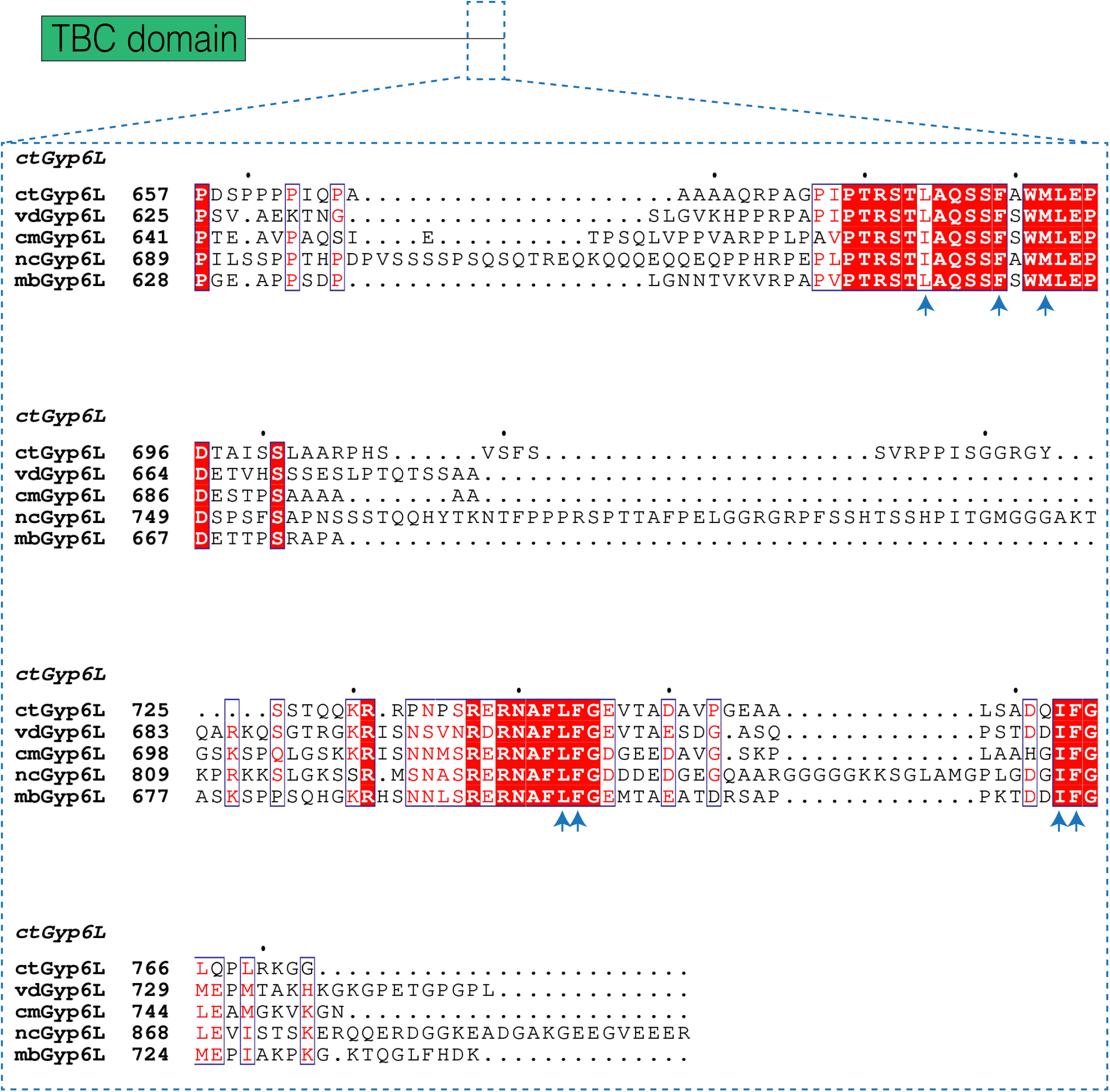
Sequence alignment of the Gyp6 C-terminal tail across the Pezizomycotina subdivision. ct, Chaetomium thermophilum; vd, Verticillium dahlia; cm, Cordyceps militaris; nc, Neurospora crassa; mb, Metarhizium brunneum. Blue arrow indicates the key residues involved in binding to Retromer.

**Supplementary Figure 12.**
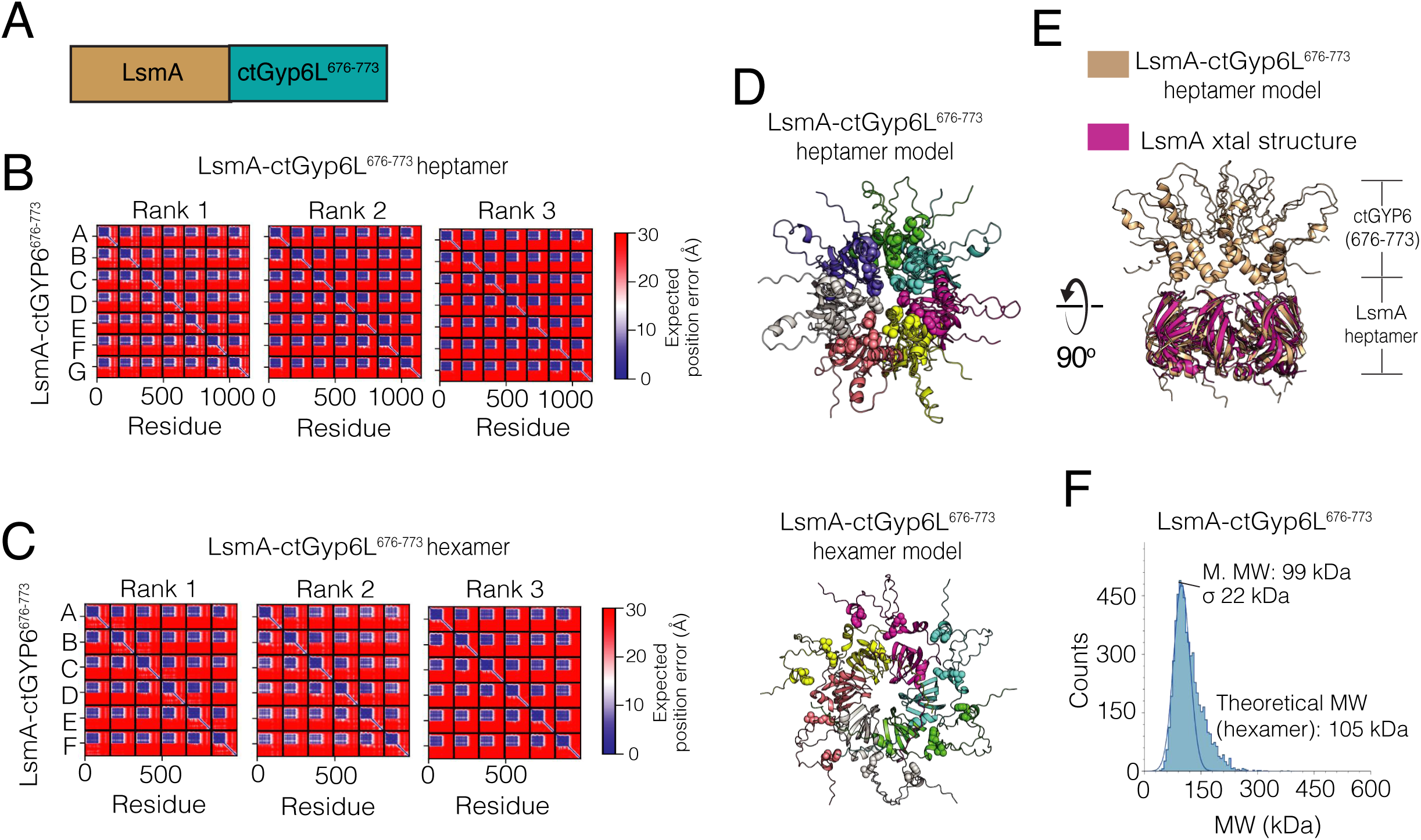
Design of LsmA-ctGyp6L^676-773^ chimeric construct to enhance ctRetromer interaction. (A) Schematic diagram showing LsmA-ctGyp6L^676-773^ chimera construct was designed by fusing ctGyp6L^676-773^ to the C-terminus of full-length LsmA. (B) PAE plot showing the top-ranked AF2 models of LsmA-ctGyp6L^676-773^ forming heptamer and (C) hexamer. (D) Cartoon representation of the top-ranked LsmA-ctGyp6L^676-773^ heptamer (top) and hexamer (bottom). For clarity, the key LF motifs in ctGyp6L_676-77_ are shown in sphere. (E) Superimposed view of LsmA-ctGyp6L^676-773^ heptamer model with the previously determined crystal structure of LsmA (PDB ID: 1I81) showing that addition of ctGyp6L^676-773^ does not affect LsmA oligomerization. (F) Mass photometry of LsmA-ctGyp6L^676-773^ showing a size corresponds to the hexamer.

---

**Table S1.**
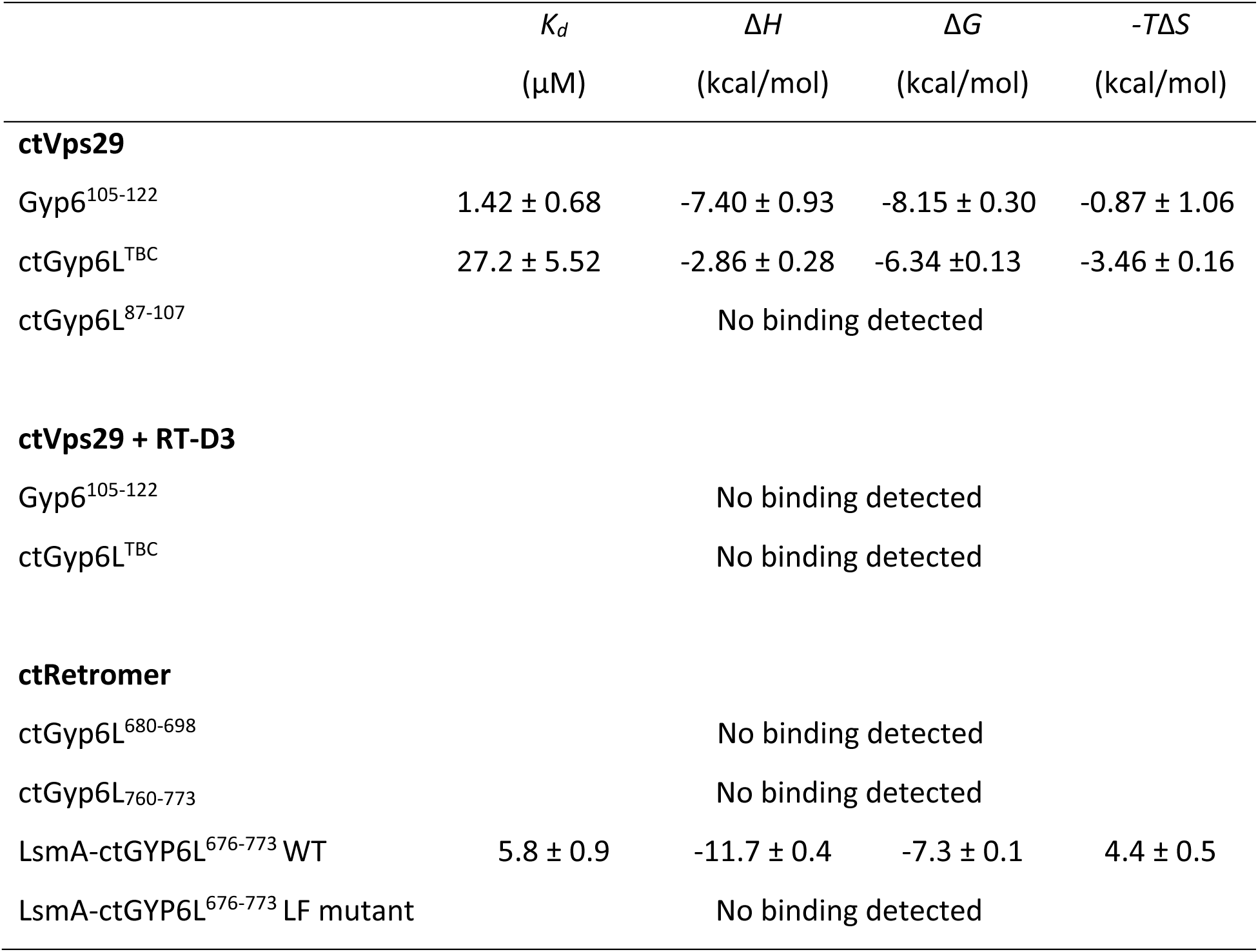
Thermodynamic parameters for the binding of Retromer and Gyp6.

| | $K_d$<br>(μM) | $\Delta H$<br>(kcal/mol) | $\Delta G$<br>(kcal/mol) | $-T\Delta S$<br>(kcal/mol) |
| --- | --- | --- | --- | --- |
| **ctVps29** |  |  |  |  |
| Gyp6<sup>105-122</sup> | 1.42 ± 0.68 | -7.40 ± 0.93 | -8.15 ± 0.30 | -0.87 ± 1.06 |
| ctGyp6L<sup>TBC</sup> | 27.2 ± 5.52 | -2.86 ± 0.28 | -6.34 ± 0.13 | -3.46 ± 0.16 |
| ctGyp6L<sup>87-107</sup> | No binding detected |  |  |  |
| **ctVps29 + RT-D3** |  |  |  |  |
| Gyp6<sup>105-122</sup> | No binding detected |  |  |  |
| ctGyp6L<sup>TBC</sup> | No binding detected |  |  |  |
| **ctRetromer** |  |  |  |  |
| ctGyp6L<sup>680-698</sup> | No binding detected |  |  |  |
| ctGyp6L<sup>760-773</sup> | No binding detected |  |  |  |
| LsmA-ctGYP6L<sup>676-773</sup> WT | 5.8 ± 0.9 | -11.7 ± 0.4 | -7.3 ± 0.1 | 4.4 ± 0.5 |
| LsmA-ctGYP6L<sup>676-773</sup> LF mutant | No binding detected |  |  |  |

**Table S2.** Summary of crystallographic structure determination statistics.

| Data collection statistics | ctGyp6L<sup>TBC</sup> | mVPS29 –<br>ctGyp6L<sup>87-107</sup> | mVPS29 - Gyp6L<sup>105-122</sup> |
| --- | --- | --- | --- |
| PDB ID | 9Z83 | 9Z7E | 9Z7D |
| Space group | P 1 21 1 | C 2 2 21 | P 1 21 1 |
| Resolution (Å) | 46.71 – 2.10<br>(2.16 – 2.10) | 38.12 – 2.13<br>(2.19 – 2.13) | 45.86 – 1.90<br>(1.95 – 1.90) |
| a, b, c (Å) | 47.93 128.42 64.50 | 44.40 72.12 114.36 | 56.86 43.35 77.31 |
| α, β, γ (°) | 90.00 102.92 90.00 | 90.00, 90.00, 90.00 | 90.00, 90.13 90.00 |
| Total observations | 301,587 (19,626) | 94,146 (7,522) | 96,263 (5,870) |
| Unique reflections | 44,048 (3,139) | 10,611 (7522) | 29,385 (1,797) |
| Completeness (%) | 98.8 (85.9) | 99.8 (98.3) | 98.3 (90.5) |
| R<sub>merge</sub><sup>+</sup> | 0.096 (1.619) | 0.129 (1.403) | 0.119 (1.486) |
| R<sub>pim</sub><sup>*</sup> | 0.039 (0.685) | 0.046 (0.484) | 0.077 (0.950) |
| CC1/2 | 0.999 (0.562) | 0.998 (0.627) | 0.994 (0.278) |
| <I/σ(I)> | 10.0 (0.8) | 9.9 (1.1) | 6.6 (1.2) |
| Multiplicity | 6.8 (6.3) | 8.9 (9.1) | 3.3 (3.3) |
| Molecule/asym | 2 | 1 | 2 |
| **Refinement statistics** |  |  |  |
| R<sub>work</sub>/R<sub>free</sub> (%) <sup>¶</sup> | 0.1888/0.2294 | 0.1899/0.2431 | 0.2636/0.2506 |
| No. protein atoms | 5,425 | 1,496 | 3,098 |
| Waters | 133 | 51 | 125 |
| Average B (Å<sup>2</sup>)<sup>^</sup> | 51.13 | 48.37 | 33.25 |
| Protein | 51.04 | 48.29 | 33.35 |
| Water | 50.35 | 50.52 | 30.26 |
| rmsd bonds (Å) | 0.008 | 0.008 | 0.009 |
| rmsd angles (°) | 1.146 | 0.99 | 1.55 |
| Ramachandran plot: |  |  |  |
| Favored/outliers (%) | 96.63/0.31 | 96.13/0.0 | 97.06/1.07 |

Values in parentheses refer to the highest resolution shell. <sup>+</sup>R<sub>merge</sub> = $\sum |I - \langle I \rangle| / \sum \langle I \rangle$ , where *I* is the intensity of each individual reflection. <sup>\*</sup>R<sub>pim</sub> indicates all I<sup>+</sup> & I<sup>-</sup>. <sup>¶</sup>R<sub>work</sub> = $\sum h |F_o - F_c| / \sum |F_o|$ , where *F<sub>o</sub>* and *F<sub>c</sub>* are the observed and calculated structure-factor amplitudes for each reflection *h*. <sup>#</sup>R<sub>free</sub> was calculated with 10% of the diffraction data selected randomly and excluded from refinement. <sup>^</sup>Calculated using Baverage.

**Table S3.** Yeast strains used in the work.

| Number | Strain | Genotype | Ref. |
| --- | --- | --- | --- |
| | BJ3505 | MATα pep4::HIS3 prb1-Δ 1.6R lys2-208 trp1-Δ101 ura3-52 gal2 can | Lab |
| | SEY6210 | MATα leu2-3,112 ura3-52 his3-Δ200 trp-Δ901 suc2-Δ9 lys2-801 GAL | Lab |
| CA239 | BJ3505 | ura3::pRS406 TetOffprom-TurboID-V5-Ypt7term | This study |
| CA240 | BJ3505 | ypt7Δ::KAN ura3::pRS406 TetOffprom-TurboID-V5-Ypt7-Ypt7term | This study |
| CA241 | BJ3505 | ypt7Δ::KAN ura3::pRS406 TetOffprom-TurboID-V5-ypt7<sup>Q68L</sup>-Ypt7term | This study |
| CA242 | BJ3505 | ypt7Δ::KAN ura3::pRS406 TetOffprom-TurboID-V5-ypt7<sup>T22N</sup>-Ypt7term | This study |
| CA260 | SEY6210 | Gyp6-mNeonGreen Vps35-yomCherry | This study |
| CA144 | SEY6210 | Vps10-mNeonGreen | This study |
| CA145 | SEY6210 | Vps10-mNeonGreen gyp3Δ | This study |
| CA220 | SEY6210 | Vps10-mNeonGreen gyp6Δ | This study |
| CA170 | SEY6210 | Vps10-mNeonGreen gyp7Δ | This study |
| CA221 | SEY6210 | Vps10-mNeonGreen gyp3Δ gyp6Δ | This study |
| CA223 | SEY6210 | Vps10-mNeonGreen gyp3Δ gyp6Δ gyp7Δ | This study |
| CA245 | SEY6210 | Vps10-mNeonGreen Vps35-yomCherry | This study |
| CA247 | SEY6210 | Vps10-mNeonGreen Vps35-yomCherry gyp3Δ gyp6Δ | This study |
| CA394 | SEY6210 | Vps35-mNeonGreen yomCherry-Ypt7 | This study |
| CA395 | SEY6210 | Vps35-mNeonGreen yomCherry- ypt7<sup>Q68L</sup> | This study |
| CA396 | SEY6210 | Vps35-mNeonGreen yomCherry-Ypt7 gyp3Δ gyp6Δ | This study |
| CA397 | SEY6210 | Vps35-mNeonGreen yomCherry- ypt7<sup>Q68L</sup> gyp3Δ gyp6Δ | This study |
| CA263 | SEY6210 | Vps10-mNeonGreen ypt7Δ ura3::pRS406 Ypt7prom- ypt7<sup>Q68L</sup>-Ypt7term gyp6Δ | This study |
| CA280 | SEY6210 | Vps10-mNeonGreen ypt7Δ ura3::pRS406 Ypt7prom- ypt7<sup>Q68L</sup>-Ypt7term gyp3Δ | This study |
| CA282 | SEY6210 | Vps10-mNeonGreen ypt7Δ ura3::pRS406 Ypt7prom- ypt7<sup>Q68L</sup>-Ypt7term gyp3Δ gyp6Δ | This study |
| CA288 | SEY6210 | Vps10-mNeonGreen ypt7Δ ura3::pRS406 Ypt7prom- ypt7<sup>Q68L</sup>- Ypt7term gyp3Δ gyp6Δ<br>leu2::pRS305-Gyp6prm-Gyp6-V5-CYC1term | This study |
| CA289 | SEY6210 | Vps10-mNeonGreen ypt7Δ ura3::pRS406 Ypt7prom- ypt7<sup>Q68L</sup>-Ypt7term gyp3Δ gyp6Δ<br>leu2::pRS305-Gyp6prm-Gyp6 R155A-V5-CYC1term | This study |
| CA314 | SEY6210 | Vps10-mNeonGreen ypt7Δ ura3::pRS406 Ypt7prom- ypt7<sup>Q68L</sup>-Ypt7term gyp3Δ gyp6Δ<br>leu2::pRS305-Gyp6prm-Gyp6 Δ415-458-V5-CYC1term | This study |
| CA323 | SEY6210 | Vps10-mNeonGreen ypt7Δ ura3::pRS406 Ypt7prom- ypt7<sup>Q68L</sup>-Ypt7term gyp3Δ gyp6Δ<br>leu2::pRS305-Gyp6prm-Gyp6 R108GL109GT110RV112GP119GL120A, Δ415-458-V5-CYC1term | This study |
| CA224 | SEY6210 | Vps10-mNeonGreen vps35Δ | This study |
| CA302 | SEY6210 | Vps10-mNeonGreen Vps35-yomCherry ypt7Δ ura3::pRS406 Ypt7prom- ypt7<sup>Q68L</sup>-Ypt7term | This study |
| CA303 | SEY6210 | Vps10-mNeonGreen Vps35-yomCherry ypt7Δ ura3::pRS406 Ypt7prom- ypt7<sup>Q68L</sup>-Ypt7term gyp3Δ gyp6Δ | This study |
| CA304 | SEY6210 | Vps10-mNeonGreen Vps35-yomCherry ypt7Δ ura3::pRS406 Ypt7prom- ypt7<sup>Q68L</sup>-Ypt7term gyp3Δ gyp6Δ<br>leu2::pRS305-Gyp6prm-Gyp6-V5-CYC1term | This study |
| CA305 | SEY6210 | Vps10-mNeonGreen Vps35-yomCherry ypt7Δ ura3::pRS406 Ypt7prom- ypt7<sup>Q68L</sup>-Ypt7term gyp3Δ gyp6Δ<br>leu2::pRS305-Gyp6prm-Gyp6 R155A-V5-CYC1term | This study |
| CA316 | SEY6210 | Vps10-mNeonGreen Vps35-yomCherry ypt7Δ ura3::pRS406 Ypt7prom- ypt7<sup>Q68L</sup>-Ypt7term gyp3Δ gyp6Δ<br>leu2::pRS305-Gyp6prm-Gyp6 Δ415-458-V5-CYC1term | This study |
| CA324 | SEY6210 | Vps10-mNeonGreen Vps35-yomCherry ypt7Δ ura3::pRS406 Ypt7prom- ypt7<sup>Q68L</sup>-Ypt7term gyp3Δ gyp6Δ<br>leu2::pRS305-Gyp6prm-Gyp6 R108GL109GT110RV112GP119GL120A, Δ415-458-V5-CYC1term | This study |

**Table S4:** Plasmids used in the work. Plasmids containing the sequence of the guide RNA for CrisperCas9 applications were produced with different auxotrophic markers, according to the strain.

| Number | Plasmid Description | Reference |
| --- | --- | --- |
| CA148 | pRS406 Ypt7prom-Ypt7-Ypt7term | This study |
| CA149 | pRS406 TetO-TurboID-Ypt7-Ypt7term | This study |
| CA150 | pRS406 TetO-TurboID-Ypt7 Q68L-Ypt7term | This study |
| CA151 | pRS406 TetO-TurboID-Ypt7 T22N-Ypt7term | This study |
| CA153 | pRS406 TetO-TurboID-Ypt7term | This study |
| CA162 | pRS305 Gyp6prom-Gyp6-V5-CYC1term | This study |
| CA163 | pRS305 Gyp6prom-Gyp6 R155A-V5-CYC1term | This study |
| CA167 | pRS406 Ypt7prom-Ypt7 Q68L-Ypt7term | This study |
| CA169 | pRS305 Gyp6prom-Gyp6 R108GL109GT110RV112GP119GL120A-V5-CYC1term | This study |
| CA176 | pRS305 Gyp6prom-Gyp6 Δ415-458-V5-CYC1term | This study |
| CA179 | pRS305 Gyp6prom-Gyp6 R108GL109GT110RV112GP119GL120A, Δ415-458-V5-CYC1term | This study |
| pSP473 | pRS423 *prGPD-cas9 his3* | Serge Pelet |
| pSP474 | pRS423 *prGPD-cas9 trp1* | Serge Pelet |
| pSP475 | pRS425 *prGPD-cas9 leu2* | Serge Pelet |
| pSP476 | pRS426 *prGPD-cas9 ura3* | Serge Pelet |

**Table S5:**
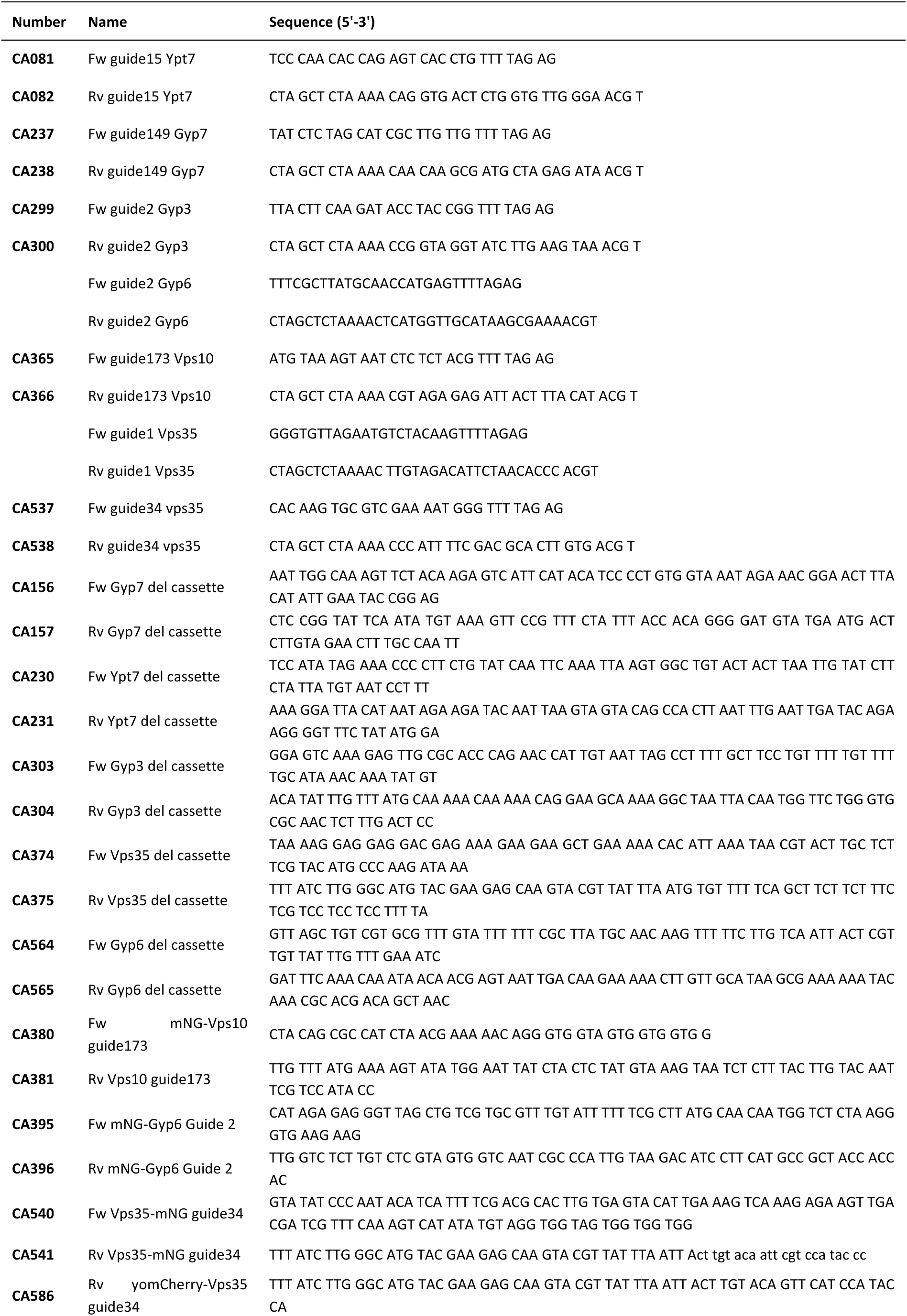
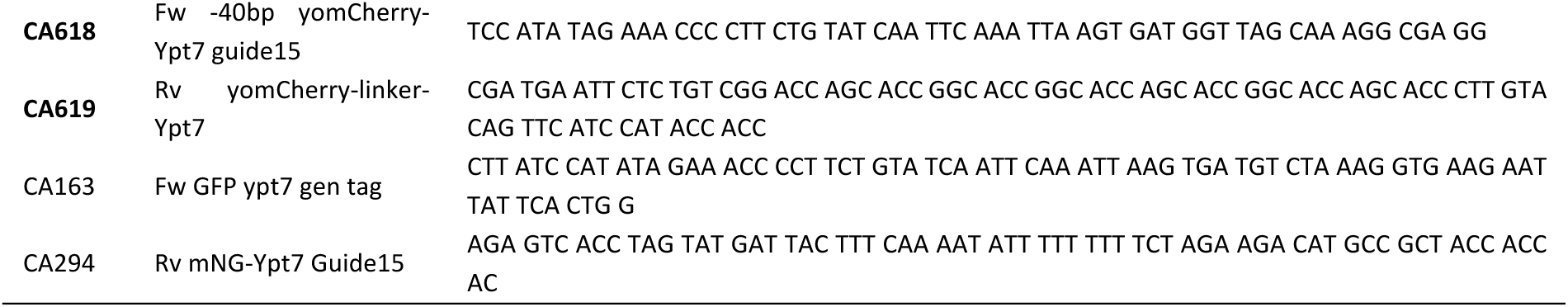
Primers used in the work.

## Notes

### Competing Interest Statement

The authors have declared no competing interest.

### Summary of Updates

Numerous figure labels in the original version had been compromised, probably due to compression artefacts. The updated version corrects this

